# Incipient local adaptation in a fungus: evolution of heavy metal tolerance through allelic and copy-number variation

**DOI:** 10.1101/832089

**Authors:** Anna L. Bazzicalupo, Joske Ruytinx, Yi-Hong Ke, Laura Coninx, Jan V. Colpaert, Nhu H. Nguyen, Rytas Vilgalys, Sara Branco

## Abstract

Human-altered environments can shape the evolution of organisms. Fungi are no exception, though little is known about how they withstand anthropogenic pollution. Here, we document incipient polygenic local adaptation in the mycorrhizal fungus *Suillus luteus* driven by recent soil heavy metal contamination. Genome scans across individuals from recently polluted and nearby unpolluted soils in Belgium revealed no evidence of population structure but detected allelic divergence and gene copy number variation in genes involved in metal exclusion, storage, immobilization, and reactive oxygen species detoxification. Standing genetic variation included multiple alleles of small effects contributing to heavy metal tolerance, suggesting the existence of different strategies to withstand contamination. These variants were shared across the whole population but under selection in isolates exposed to pollution. Together, our results point to *S. luteus* undergoing the initial steps of adaptive divergence and contribute to understanding the processes underlying local adaptation under strong environmental selection.

## Introduction

Understanding how environments shape evolutionary change is a long-standing topic in biology^1–5^. Environmental shifts are particularly well-known for affecting the evolution of organisms and leading to local adaptation, with individuals displaying higher fitness in their original environment^6^. Novel environmental conditions can offer strong selection that acts on phenotypes and respective genotypes^7^. Such selection can lead to the evolution of local adaptation, with specific genotypes and phenotypes allowing establishment in the new environments. Full documentation of the processes and mechanisms of local adaptation has been challenging, but the increasing ability to detect and test the genomic regions under selection in a wide range of systems has allowed unprecedented advancements in this field^8,9,10,11^.

Local adaptation imprints genomic signatures that are easiest to detect when selection is strong and acts on few genomic regions of large effects. In this scenario, a small number of genetic variants dramatically increase fitness under specific environments and quickly spread through the population. Such process creates a distinct genetic signal, with the few genomic targets of selection highly differentiated and deviating from the remaining genome^12,13^. Examples of this phenomenon include the evolution of pesticide resistance, such as in the insects *Anopheles*^14,15^ and *Drosophila*^16^, and fungicide resistance in the fungal pathogen *Zymoseptoria*^17^. However, selection associated with environmental shifts can instead target complex traits that involve a multitude of genes and genetic pathways^18,19^. Such polygenic adaptation has been described in tabletop corals^10^, maize^9^ and the fungus *Neurospora*^20^ evolving thermal tolerance under changing climatic conditions. Detecting complex trait adaptation is challenging, as it translates into low differentiation across many sites that are difficult to distinguish from the background genomic signal. The ability to detect polygenic selection decreases with the number of alleles contributing to the trait, as a multitude of alleles under selection results in both a weakened signal of selection^21^ and very low divergence across the genome^18^. Furthermore, alleles of small effects are predicted to show genetic redundancy and standing variation^18^. This means multiple loci independently confer adaptive phenotypes and are also present at low frequencies in populations not under selection. Another important expectation of small effect alleles is their susceptibility to gene flow between locally adapted and nearby non-adapted populations, which can lead to the loss of adaptive alleles if selection is not sufficiently strong^22–24^. Here, we investigate the genomic signatures of heavy metal adaptation in *Suillus luteus*, a pine mutualistic fungus. Our goal was to identify whether this system followed a ‘few alleles of large effects’ or a ‘many alleles of small effects’ pattern and clarify the role of standing genetic variation in the adaptive process.

Human-induced environmental shifts, such as pollution and climate change, provide unique opportunities to study the process of local adaptation and its genetic signatures. These shifts tend to be associated with the evolution of complex traits^25–27^ and polygenic adaptation^5,10,28–30^, making it challenging to identify relevant adaptive alleles. There are however good examples of the genetic basis of polygenic adaptation to anthropogenic activity, involving multiple physiological traits and gene pathways^30,29^. Anthropogenic heavy metal pollution in soils can also impose serious environmental shifts and offer strong selection leading to change in complex traits, as it tends to include a multitude of stressors that affect different physiological responses and is easily lethal^31,32^. Furthermore, even though high concentrations of heavy metals are always toxic due to oxidative stress caused by the production of reactive oxygen species (ROS), some heavy metals, such as zinc, iron, and copper, are essential micronutrients and needed in small amounts for proper cell functioning^33,34^. These metals cannot therefore be completely excluded from cells and tight regulatory systems ensure balance between heavy metal detrimental excess and deficiency^33,35,36^. Metal homeostasis systems are based on four main concomitant strategies: metal exclusion, storage, immobilization, and ROS detoxification^35,37^. Metal exclusion and storage is achieved through transmembrane transporter proteins that are responsible for moving metal ions across membranes and import or extrude metal ions to/from the cell or into membrane-bound vesicles or vacuoles. Metal immobilization is carried out by chelating agents that bind to and inactivate metal ions. Finally, ROS detoxification occurs when antioxidants act to reduce oxidative stress caused by ROS produced as a response to high cellular metal content. Given that metal exclusion, storage, immobilization, and ROS detoxification are based on the action of many different genes and functions, the genomic signatures of heavy metal adaptation are likely polygenic, with low differentiation in a large number of alleles with rather small effects in numerous metal regulatory pathways. In addition, along with allelic variance, heavy metal tolerance is also known to be achieved through differences in gene copy number, with tolerant and non-tolerant individuals carrying different numbers of copies of specific adaptive alleles^38^.

Despite the ubiquitous presence of fungi in soil and known examples of fungal heavy metal tolerance^39–42^, very little is known on the genetic basis of fungal metal pollution adaptation. Here we aim to fill this gap by investigating the evolution of heavy metal tolerance in *Suillus luteus* (Boletales, Basidiomycota), a mycorrhizal soil fungus associated with pine trees (Fig. 1). Species of the genus *Suillus* are known to be tolerant to heavy metal contaminated soils^43,44^ and previous research showed pronounced heavy metal tolerance in *S. luteus*. Specifically, isolates from heavy metal polluted regions in Belgium were found more likely to be tolerant to high levels of cadmium and/or zinc, while isolates from nearby non-polluted areas are less likely to survive in high metal concentrations^41,45–50^. Contrary to non-tolerant isolates, metal-tolerant *S. luteus* grow well at high cadmium and zinc concentrations while accumulating surprisingly low amounts of metals in their tissue, suggesting stark differences in metal homeostasis across metal-tolerant and non-tolerant isolates. Targeted genetic studies in the same Belgian *S. luteus* isolates identified four transporter genes putatively involved in zinc homeostasis (*SlZRT1, SlZRT2, SlZnT1*, and *SlZnT2*) ^45,51,52,53^, suggesting these genes play an important role in tolerating contaminated sites. Survival on soils with several contaminants such as zinc and cadmium also suggests *S. luteus* has multiple strategies to maintain heavy metal homeostasis under high metal conditions.

**Figure 1.**
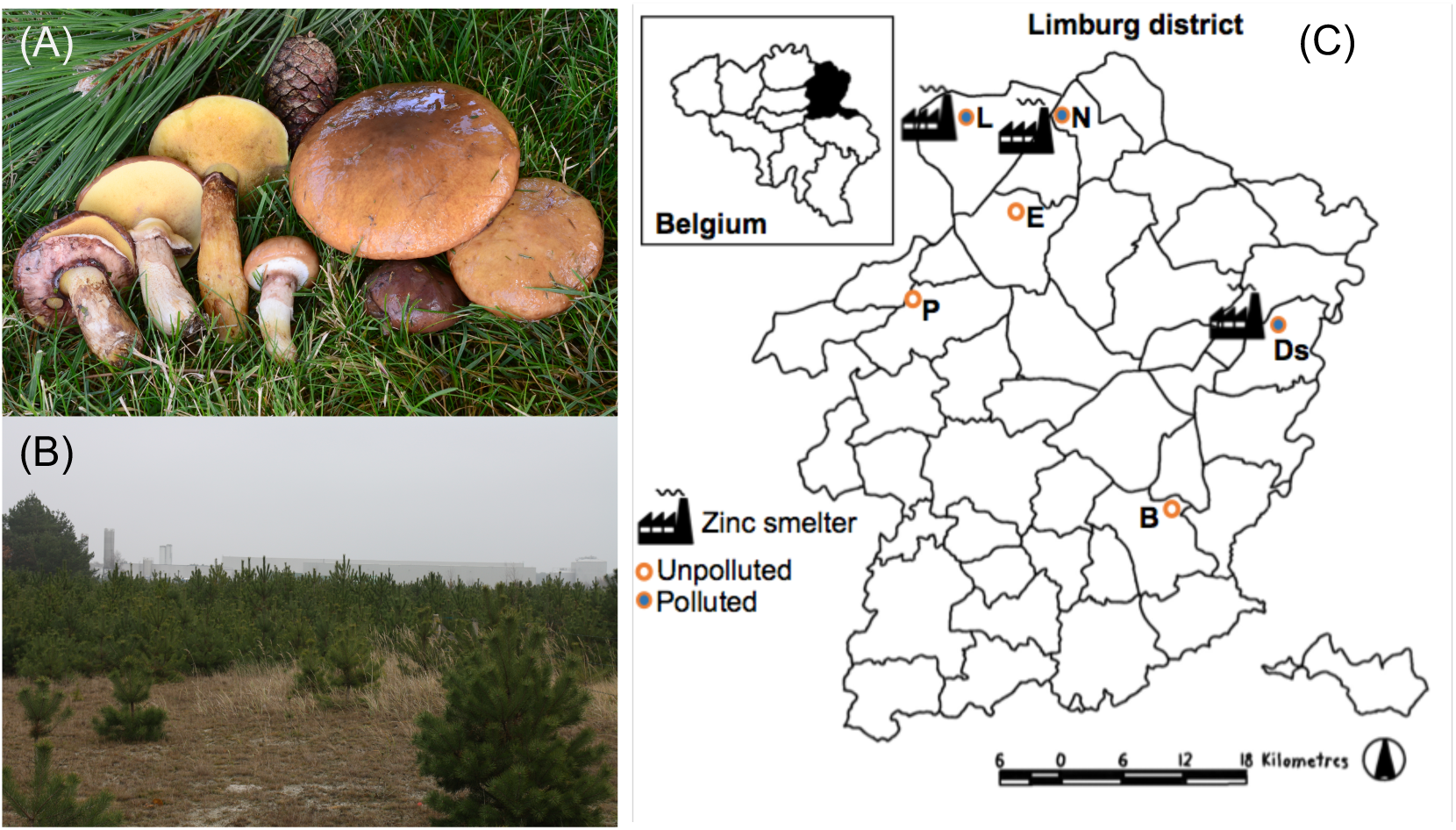
(A) *Suillus luteus*, a mycorrhizal fungus associated with pines (photo by Noah Siegel). (B) Lommel, one of the sampled polluted sites (photo by Joske Ruytinx). (C) Sampled localities in Belgium (Limburg district): three heavy metal polluted (full circles) and three unpolluted sites (empty circles). B-Bilzen, Ds-Dilsen-Stokkem, E-Eksel, L-Lommel, N-Neerpelt, P-Paal. (Smelter icon http://chittagongit.com/icon/factory-icon-transparent-6.html).

We investigated the evolution of heavy metal adaptation in *S. luteus* by comparing whole genomes of isolates from polluted and adjacent non-polluted sites in Belgium (Fig. 1). Based on the striking observed metal tolerance differences across isolates from such a small geographic region^48,54^, we hypothesized that (1) individuals from the two soil types have overall low genomic differentiation, (2) heavy metal tolerance stems from differentiation in many alleles of small effects present across the whole population and (3) differentiation is highest in genes involved in metal homeostasis. We found no population structure and very low divergence across isolates from the two soil types, consistent with ongoing gene flow and a single outcrossing population. We also detected signatures of polygenic adaptation, with many loci and gene pathways showing both allelic divergence and copy-number variation. As expected, many differentiated genes were involved in metal exclusion, storage, immobilization, and ROS detoxification functions. These results point to incipient local adaptation, with standing genetic variation including alleles advantageous in polluted soils that are under positive selection in *S. luteus* isolates exposed to high heavy metal pollution.

## Results

### Absence of *Suillus luteus* population structure across polluted and unpolluted sites

Whole genome analyses provided no evidence of population structure across sampled sites or soil type (Fig. 2). This result confirms previous fragment analyses^48^ and suggests ongoing gene flow between isolates from polluted and non-polluted sites. Analyses of ~1.6 million high quality single-nucleotide polymorphisms (SNPs) obtained by comparing 38 *S. luteus* whole genomes (see Table S1 for details) revealed one population as the most likely explanation of the genetic variant data (Fig. S1) and showed no isolate clustering based on site or soil pollution (Fig. 2). Four of the nine isolates from the Lommel site did not cluster with the main group of isolates in this analysis (Fig. 2), while the other five isolates grouped with the rest of the isolates studied. Such disparity may reflect introgression of genetic material from nearby populations, however we are unable to test this hypothesis. Window summary statistics of genomic variation were also consistent with a single population and ongoing gene flow across all sampled sites. We found comparable estimates of nucleotide diversity (π) for all samples (0.00649), for isolates from polluted sites (0.00645), and isolates from unpolluted sites (0.00650), and also recovered a negative value for Tajima’s D (=−0.29696), suggestive of a past bottleneck.

**Figure 2.**
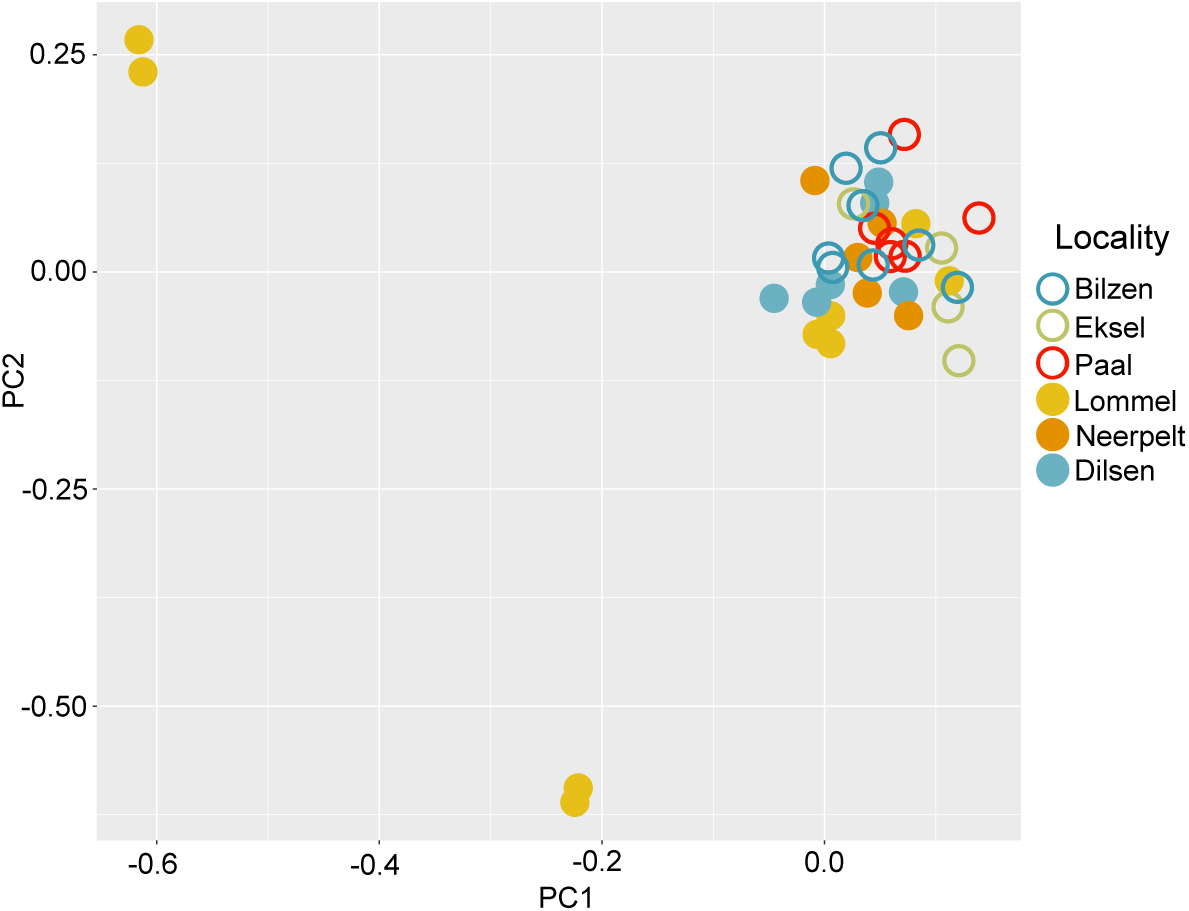
Absence of population structure across *S. luteus* isolates. Principal Components Analysis based on ~1.6 million SNPs from genomes sampled on heavy metal polluted (full circles) and unpolluted (empty circles) sites in Belgium.

### Low overall allelic divergence, with most diverged regions identified as transmembrane transporters

We detected low genomic divergence across whole genomes of isolates from polluted and non-polluted soils. This result is concordant with one panmictic population and ongoing gene flow. Genomic differentiation included many alleles scattered across the genome and was enriched for transmembrane transporters. We measured genomic divergence by calculating allele frequency differences (F_ST_) and the average number of nucleotide differences (d_XY_) per 5kb windows and found both to be overall low (average F_ST_ = 0.003, Fig. 3A, average d_XY_=0.001, Fig. S2). Gene ontology (GO) enrichment analyses on the top 5% differentiated genomic regions based on F_ST_ and d_XY_ identified the most representative gene categories putatively differentiating samples from polluted and unpolluted soils. We found the top 5% variants included genes with functions related to heavy metal tolerance, including metal exclusion, storage, immobilization, and ROS detoxification. Close to 20% of the most highly F_ST_ differentiated genomic regions were significantly enriched for transport functions, important in the exclusion and storage of heavy metals outside the cytoplasm^34,36,55–57^ (Fig. 3B, Table S2). The top 5% d_XY_ regions were differently enriched, with ~40% represented by GO term ‘protein kinase activity’ (Fig. S3). The d_XY_ and F_ST_ enrichment discrepancy might be explained by the fact that d_XY_ considers allelic differences but not their frequency^58^ and could fail to capture genomic regions under selection in a system with gene flow and standing genetic variation (see below). Notably, none of the genes identified in previous studies (*SlZRT1*, *SlZRT2*, *SlZnT1* and *SlZnT2*) with a role in *S. luteus* zinc tolerance^51,52^ were in the top 5% diverged windows, however, the F_ST_ value for *SlZnT2* did fall within the top 10% (*SlZnT2* F_ST_ = 0.0200982).

**Figure 3.**
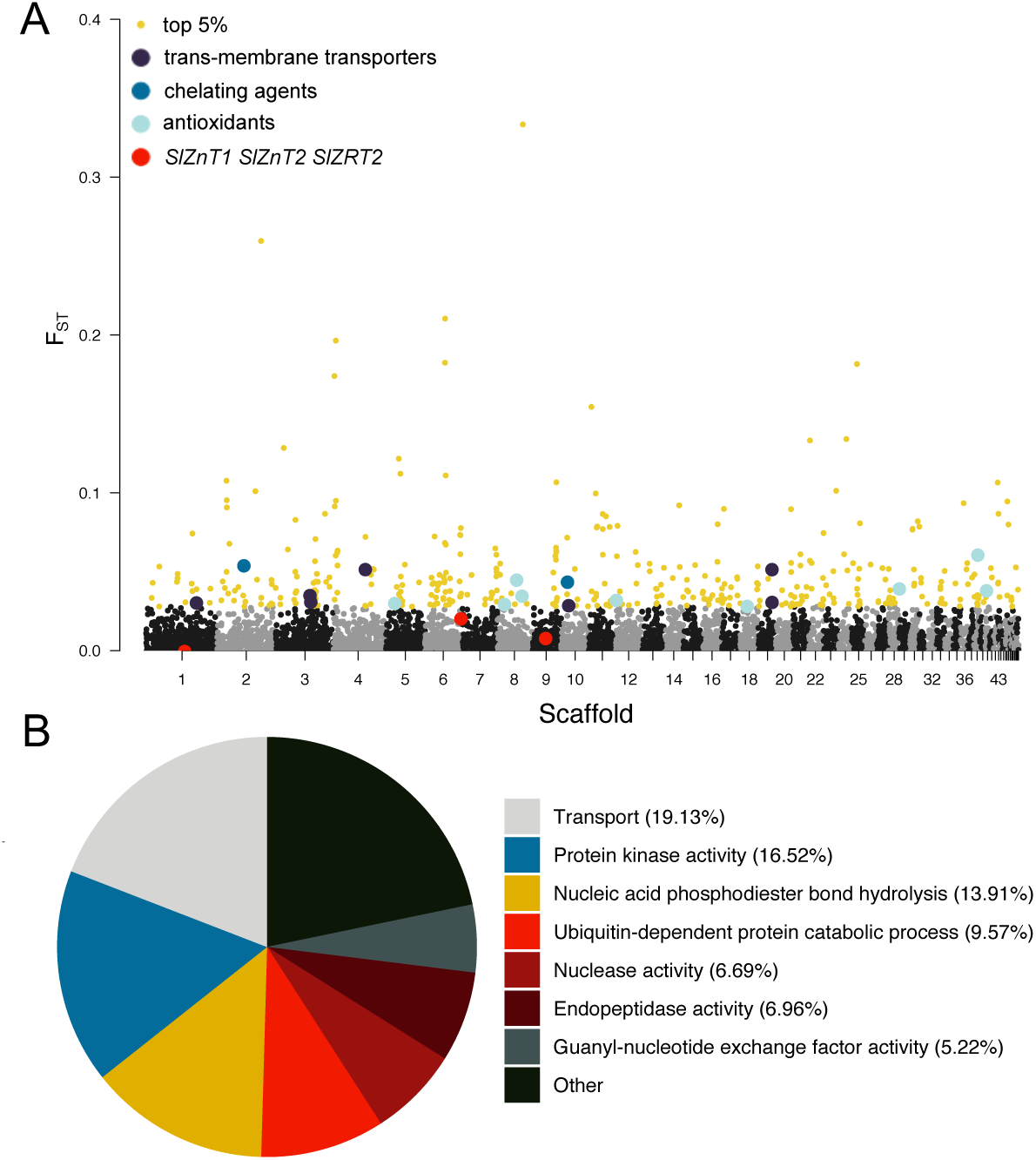
(a) Average F_ST_ values across 5-Kb genomic windows between *Suillus luteus* from polluted and unpolluted soils. Yellow dots show the top 5% F_ST_ values. Loci involved with transport are shown in dark purple, chelators in blue and antioxidants in light blue (see Table 1 for details). The previously studied zinc transporters (*SlZnT1, SlRT2*, and *SlZnT2*,the F_ST_ for the window containing *SlRT1* is just below zero) are depicted in red. (b) Gene ontology (GO) enriched terms in the top 5% FST values, with respective percentages.

### No genetic variants were significantly associated with heavy metal polluted soil

We conducted a genome-wide association study (GWAS) to explicitly identify common variants of modest effects^59^ and failed to detect any specific genetic variants significantly associated with heavy metal pollution. We used a case/control approach that compared isolates from polluted (case) and unpolluted (control) soils and estimated their association significance to each SNP. We found no significant variants following a Bonferroni correction or a permutation test. Even though we found no significant differences (most likely as a result of our small sample size^21^), one of the variants with top p-values (p=1.54e^−7^) included six SNPs located in the promoter region of a Rab geranylgeranyltransferase (Fig. 4). This protein interacts with the membrane and is predicted to have a zinc binding site^60^, suggesting an association between the expression of this gene and the occurrence in polluted sites.

**Table 1.** Differentiated genes involved in heavy metal tolerance strategies (see Table S4 for details). We combined the metal exclusion and storage strategies as they both involve trans-membrane transporters.

| Heavy metal tolerance strategy | Category (# loci) | Predicted/potential function | Annotation |
| --- | --- | --- | --- |
| **EXCLUSION /STORAGE:** | Metal ion transporters (4) | Zinc transporter from cytosol into vesicle/vacuole<sup>34</sup> | Cation diffusion facilitator (CDF) transporter |
|  |  | Iron transporter from cytosol into vesicle/vacuole<sup>37</sup> | Iron permease (FTR1) |
|  | Transport and transfer proteins, interacting with membranes (11) | Vesicle transport<sup>64</sup> | ER-Golgi trafficking TRAPP I complex 85 kDa subunit |
|  |  | Vesicle transport<sup>65,66</sup> | Armadillo-type fold |
|  |  | Vesicle transport from Golgi<sup>67</sup> | Vesicle transport protein, Got1/SFT2-like protein |
|  |  | Vesicle transport<sup>68</sup> | Exocyst complex component Sec10-like protein |
|  |  | Vesicle transport<sup>69</sup> | OPT oligopeptide transporter protein |
|  |  | Zinc binding site<sup>60</sup> | Rab geranylgeranyltransferase |
|  |  | Inhibit Rab geranylgeranyltransferase<sup>70</sup> | Farnesyltransferase |
| **IMMOBILIZATION:** | Chelating agents within cytosol (4) | Synthesis of Nicotianamine a cadmium chelating agent in cytosol<sup>42</sup> | S-adenosyl-L-methionine-dependent methyltransferase |
|  |  | Zinc chelating in cytosol<sup>71</sup> | HIT-like protein |
|  | Chelating agents outside the cell (4) | Response to stress and chelating secreted protein<sup>37</sup> | Heat shock protein 70 family |
|  |  | Metal ion chelating secreted protein<sup>37</sup> | Fungal hydrophobin-domain containing protein |
| **ROS DETOXIFICATION:** | Oxidative stress relief (24) | Antioxidant<sup>72</sup> | Manganese/iron superoxide dismutase |
|  |  | Antioxidant<sup>73</sup> | FAD-binding |
|  |  | Antioxidant<sup>73</sup> | Ferric reductase NAD binding domain |
|  |  | Antioxidant<sup>62</sup> | Glutathione S-transferase |
|  |  | Pollution detoxification<sup>63</sup> | Cytochrome P450 |

**Figure 4.**
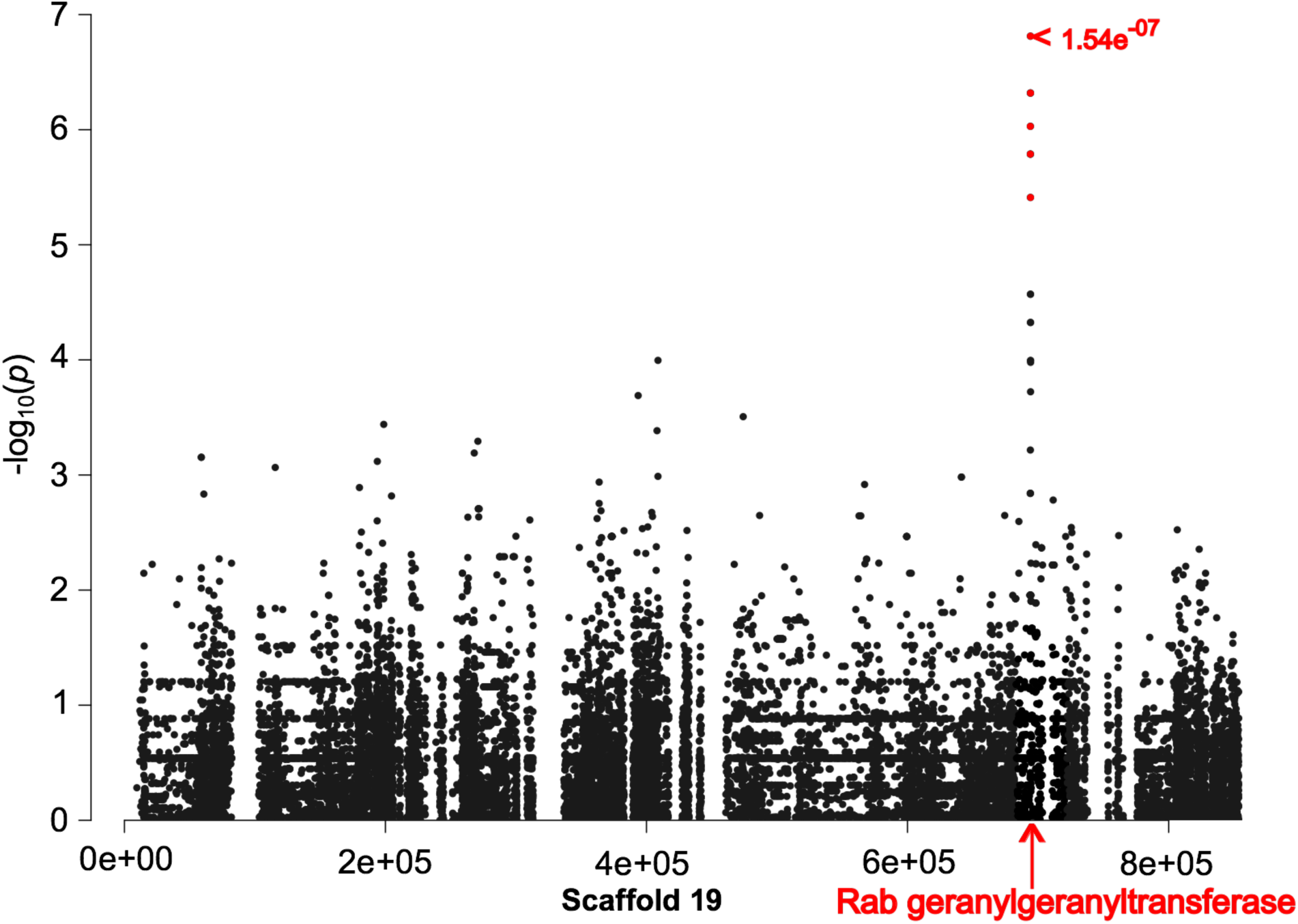
Genome wide association between SNPs and soil type. The six SNPs with highest p-values (in red, two dots are overlapping where 1.54e^−07^ is indicated, non-significant based on a Bonferroni correction where p=1e^−08^) are predicted to be located in the promoter region of gene Rab geranylgeranyltransferase, a zinc binding transferase (in red). Dots corresponding to the SNPs within the coding sequence of the gene are indicated by the bottom arrow.

### Genes with high copy-number variation are involved in transport activities

We found wide variation in gene copy-numbers across the genome, with ~40% of the most highly differentiated genes involved in transport activities (Fig. 5). This result points to many genes contributing to overall heavy metal tolerance through both gene multiplication and deletion. The top 5% copy-number variant gene ontology enrichment analysis revealed transporters as the most represented category differentiating isolates from polluted and non-polluted sites (Fig. 5B, Table S3). Highly differentiated variants also included genes with other functions related to heavy metal homeostasis (Table 1). Gene deletions and multiplications occurred on both soil types, with no clear pattern of either one associated with colonizing polluted and unpolluted soils (Fig. 6).

**Fig 5.**
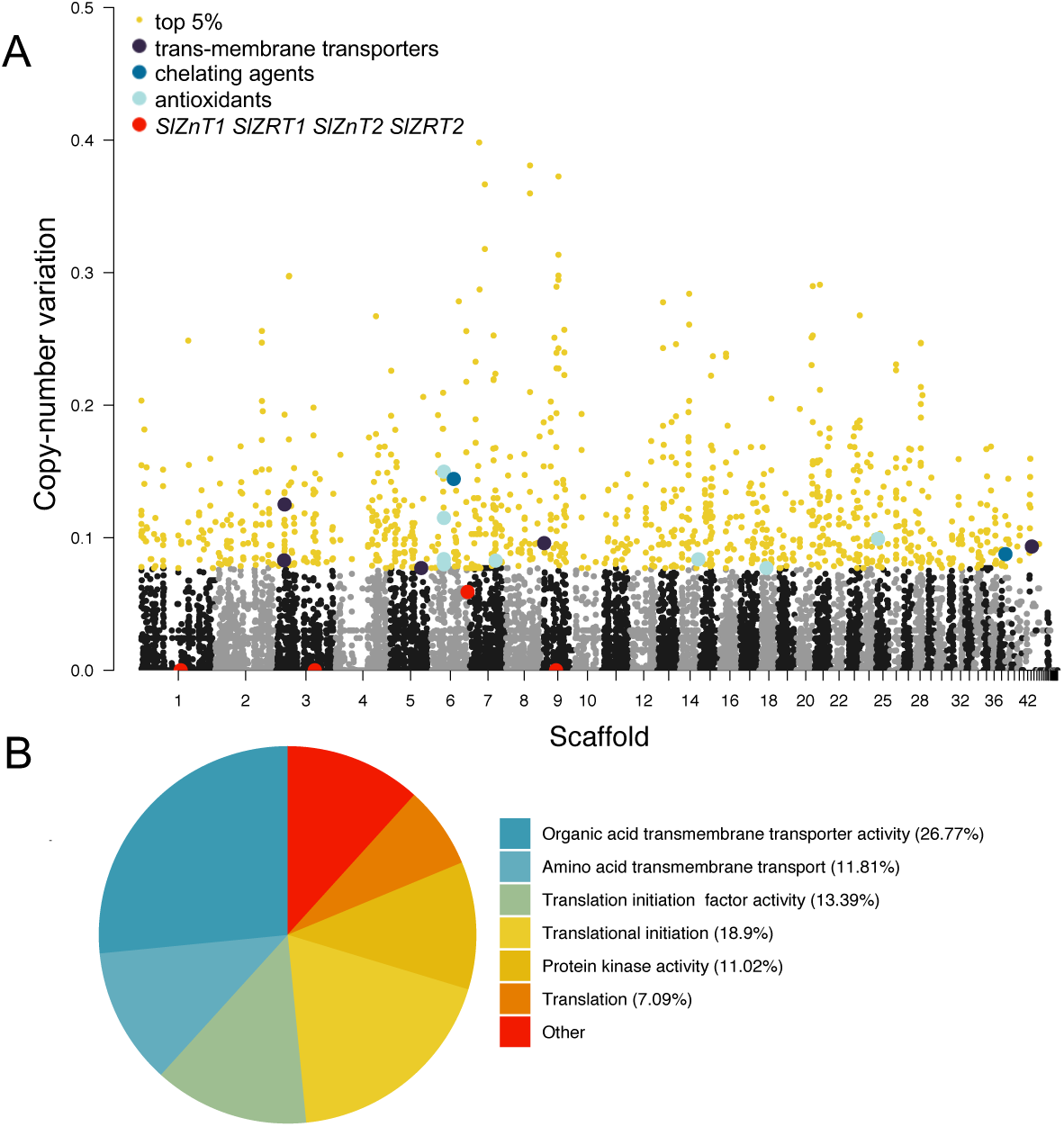
(a) Average gene copy-number variation across 5-Kb genomic windows between *Suillus luteus* from polluted and unpolluted soils. Yellow dots show the top 5% gene copy variation values. Genes involved with transport are shown in dark purple, chelators in blue and antioxidants in light blue (see Table 1 for details). The previously studied zinc transporters (*SlZnT1, SlZnT2*, *SlRT1*, *SlRT2*) are depicted in red. (b) Gene ontology (GO) enriched terms in the top 5% FST values, with respective percentages.

**Figure 6.**
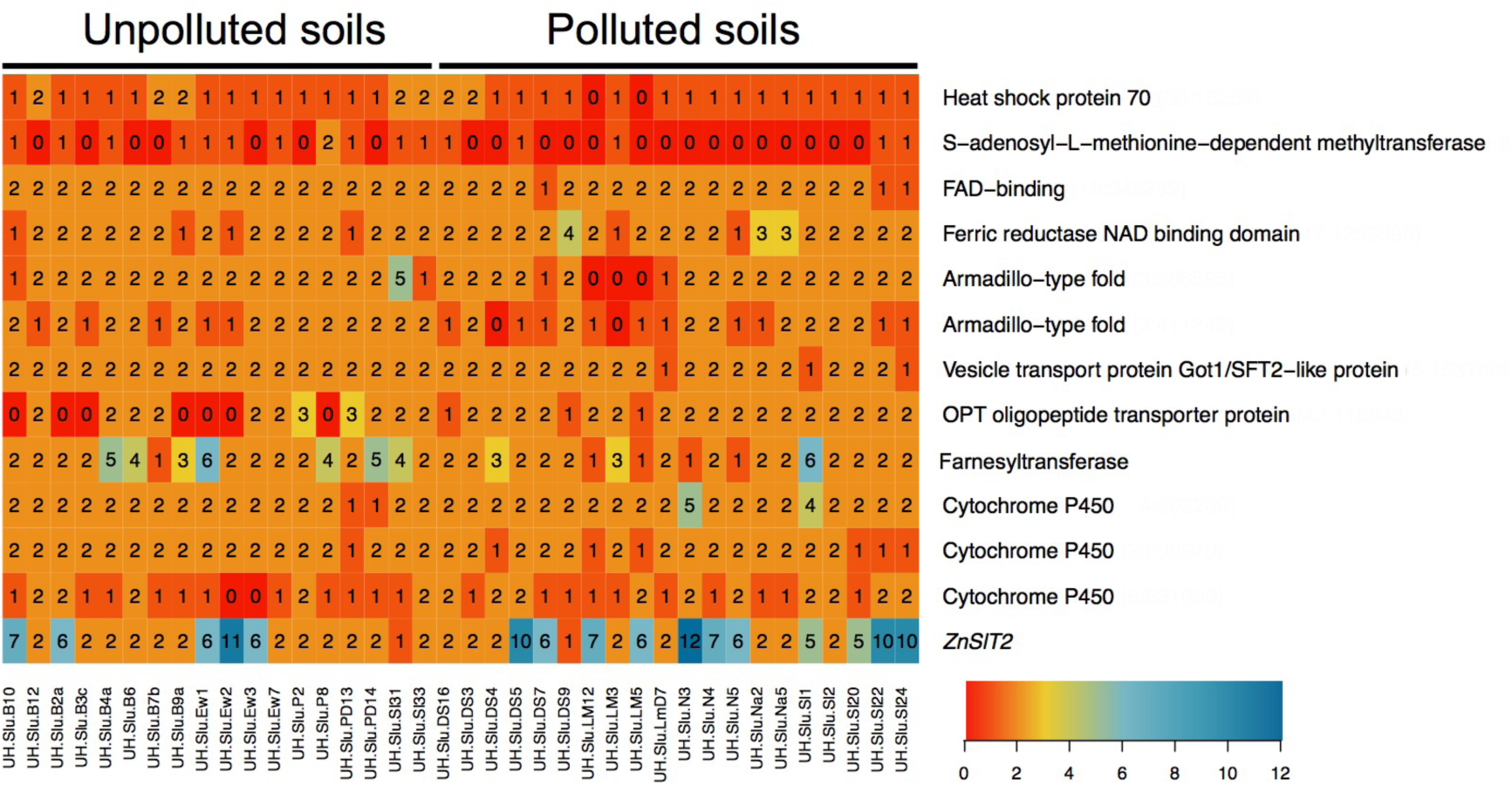
Gene copy number variation in top 5% genes involved in heavy metal tolerance. Heatmap of genes (rows) with copy number of each isolate (columns) between polluted and unpolluted localities. Lower right: gene copy number color key.

Consistent with previous findings^61^, we identified isolates from unpolluted soils with multiple copies of *SlZnT2* that are likely zinc tolerant (found in the top 10% of our copy number variation estimates and shown to be correlated with zinc tolerance^53^), as well as isolates with low number of *SlZnT2* copies (shown to be sensitive to high zinc^53^) inhabiting polluted soils (Fig. 6). These findings could derive from the existence of soil heterogeneity^54^, with localized lower levels of soil contamination that can allow growth of non-tolerant isolates. These results are also consistent with ongoing gene flow between polluted and non-polluted sites and with standing genetic variation including variants associated with metal tolerance. Based on our data we cannot determine whether current observed standing genetic variation was present prior to the polluting event or come about by the spread of tolerant genotypes from polluted areas through gene flow.

### *Suillus luteus* heavy metal tolerance candidate genes

Of the top 5% differentiated genes for allele and copy-number variants, we identified 47 gene candidates scattered across the genome putatively involved in heavy metal tolerance (Table 1 and Table S4). These candidate genes encode for transmembrane transporters involved in metal exclusion and storage, immobilization, and ROS detoxification. The large number of candidate genes associated with metal homeostasis suggests a moderate individual gene contribution to heavy metal tolerance in *S. luteus*. Fifteen candidate genes were involved in metal exclusion and storage, with four genes predicted to actively pump metal ions across membranes located either in the plasma membrane (contributing to extruding metals to the cell exterior) or vesicle and vacuole membranes (creating metal storage that can be either kept in the cell or extruded). Examples of these genes include the cation diffusion facilitator transport proteins that are predicted to aid in zinc ion homeostasis^34^ and an iron permease gene predicted to transport iron ions across membranes but also reported to interact with other metal such as zinc and cadmium^33,37^. We detected many more genes involved in membrane transport and metal exclusion and storage (Table 1), however it is unclear how exactly they contribute to heavy metal tolerance in *S. luteus*. We also found eight genes encoding for chelating agents that are responsible for immobilizing metals and preventing oxidative stress^34^, including three S-adenosyl-L-methionine-dependent methytransferases that are involved in the synthesis of the cadmium chelator nicotianamine^33,42^. In addition, we detected 24 genes involved in ROS detoxification. These included glutathione S-transferase, a key player in reducing oxidative stress caused by heavy metal toxicity^62^, and fifteen loci predicted as Cytochrome p450, a protein family known to be involved in detoxification of environmental pollutants^63^.

## Discussion

We investigated the genomes of *S. luteus* individuals from nearby heavy metal polluted and non-polluted soils and found no evidence of population structure (corroborating a previous study based on fragment analysis^48^) and high levels of shared genetic variation. We also found allelic and gene copy number differentiation spread across the genome that included many genes related to metal homeostasis. These results are consistent with incipient local adaptation, with ongoing gene flow and standing genetic variation including many loci contributing modestly to heavy metal tolerance.

Several factors are likely to contribute to the detected patterns, namely recent pollution (~150 years), the absence of barriers across polluted and unpolluted soils, and the ability of a few metal tolerant and sensitive isolates to persist in non-polluted and polluted soils^61^. Such reciprocal presence of tolerant and sensitive isolates in both soil types is likely due to both the absence of a strong trade-off between metal tolerance and survival in non-polluted soil and the occurrence of pockets of low contamination in polluted sites that allow metal sensitive isolates to persist. The *S. luteus* low overall divergence pattern of selection contrasts with other systems showing local adaptation that show clear population structure and few alleles with major effects driving divergence^14,16,17,29^. However, there are other reports of local adaptation with no population structure, low genomic divergence, and many genes of small effects. For example, local adaptation to low and high salinity in oysters was found to include a single highly outcrossing population with overall low divergence due to the lack of barriers to geneflow, and small fitness contributions from alleles determining salt tolerance^74^. Local adaptation is only expected to lead to population differentiation if selection in the new environment is severe enough to prevent the survival of genotypes lacking specific adaptive alleles^6^ and/or the absence of gene flow that homogenizes the gene pool^23^. Given the shared standing genetic variation, the absence of gene flow barriers across soil types, and the potential for heterogeneity in soil chemistry, it is very unlikely that this scenario of incipient local adaptation in *S. luteus* will evolve into population differentiation or speciation in the future.

Adaptation to high heavy metals is known to occur with both allelic divergence and gene copy number variation, with tolerant and non-tolerant individuals carrying distinct adaptive alleles^29,75^ and/or different adaptive allele copy number^38^. For example, copper tolerance in yeast is achieved by both allelic and copy number variation mutation in genes belonging to a series of enzymatic pathways, especially related to metal storage and involving vacuolar transport genes^76^. Gene copy number variation has also been documented to confer cadmium tolerance in populations of the plant *Noccaea caerulescens*^77^ and zinc and cadmium tolerance in populations of *Arabidopsis halleri*^75^. In *S. luteus*, standing genetic variation included both allelic and gene copy number variation, where variants conferring higher fitness in the new environment are present at low frequency across the entire population^78^. Although not advantageous in the original environment, these shared variants are instrumental in allowing individuals to colonize and survive in the novel environmental conditions^6^. Widespread standing genetic variation is also evident in *S. luteus* through the overall very low allelic and copy number differentiation across isolates from different soil types and adaptive alleles present in isolates across the population, as expected by theory^78^. A specific example is the gene *SlZnT2*, with high number of copies known to associate with zinc tolerance^53^ present in isolates from both polluted and unpolluted soils.

Differentiation across *S. luteus* isolates from polluted and non-polluted soils included many alleles of modest effects, with the targets of selection broadly distributed across the genome and different genetic combinations conferring ability to persist in contaminated soils. This last result corroborates a previous experimental studies showing metal-tolerant isolates display multiple distinct genotypes^53^. In addition, the zinc transporters *SlZnT1* and *SlZnT2* genes that were not included in the top 5% candidate genes under selection have been experimentally documented to be involved in zinc tolerance^45,52^, indicating that genes not primarily targeted by selection also contribute to heavy metal tolerance. Such polygenic adaptation is common and has been reported in systems with selection acting on complex traits, such as tolerance to temperature, pollutants, and serpentine soils. For example, killifish exposed to increased water temperatures^28^ and a cocktail of pollutants^79^ evolved tolerance through differentiation in many different genes associated with a multitude of physiological responses, and different genotypes conferring tolerance to the new environment^79^. Likewise, North Atlantic eels when exposed to pesticides and heavy metals, showed differentiation occurring in alleles of many genes in different pathways involved in sterol regulation^29^. Polygenic adaptation is also known in cultivated maize and was key to the success of domestication across many different environments in North America^9^. Lastly, serpentine soils are well known example of compound environmental stresses that promoted polygenic adaptation in *Arabidopsis* involving tolerance to both drought and high concentrations of heavy metals^80^. The studied *S. luteus* population is challenged by multiple heavy metals and likely also the effects of other abiotic factors (such as pH variation affecting the bioavailability of metal ions) that influence its ability to persist in polluted soils^41^. Polygenic adaptation is therefore an expected response to heavy metal pollution.

Soil heavy metal pollution is widely known for shaping the evolution of organisms^31^ and heavy metal tolerance is known to involve genes in a multitude of pathways^35,36^. As most other organisms, *Suillus luteus* metal tolerance is achieved through selection acting on many genes involved in metal homeostasis, specifically in metal exclusion, storage, immobilization and ROS detoxification strategies. Metal exclusion and storage is undertaken by transmembrane transporters that regulate the movement of metal ions across plasmatic and organelle membranes, actively excluding toxic metals from the cytoplasm. Metal immobilization occurs through the action of chelating agents. In fungi these can occur both outside and inside the cell, with for example hydrophobins and melanins binding metals to the fungal cell wall^37^ and nicotianamine as key in maintaining metal ion homeostasis within the cell^81–83^. ROS detoxification occurs when antioxidants (e.g. superoxide dismutases, ferric reductases, Cytochrome p450, and glutathione S-transferases^62,72,73^) act to reduce oxidative stress caused by ROS produced as a response to high cellular metal content, defending the cell from oxidative damage caused by heavy metals ions^79^. These strategies act together and are predicted to occur in different parts of the fungal hypha (Fig 7). The main metal tolerance strategy in *S. luteus* is metal exclusion/storage, with transporter proteins as the primary targets of selection. Transmembrane transporters are known for being crucial for heavy metal tolerance^34^ and differentiated transporters included a cation diffusion facilitator transporter and other transporters involved in metal transport also in plants and yeast, such as the ZIP (Zrt-, Irt-like Protein) protein transporter family^34,55^ that transports micronutrients^57^ across membranes to and from the cell and organelles. Notably, the four transporters previously found to be involved in zinc homeostasis were not among the most highly differentiated genes. These were two plasma membrane zinc importer genes in the ZIP (Zrt-, Irt-like Protein) family^45,51^ and two cation diffusion facilitators (CDFs) involved in zinc storage in the vacuole^52,53^. Functional assays showed that *SlZRT1* transcription is greatly reduced when *S. luteus* is exposed to toxic concentrations of zinc^45^ and heterologous expression in yeast and transcription assays of *SlZRT2* suggest this gene is involved in redistribution of zinc inside the cell^51^. Furthermore, vacuolar localization and zinc specificity of *SlZnT1* was confirmed with heterologous expression of the gene in yeast. In addition, increased zinc tolerance was found to be correlated with an increased number of *SlZnT2* copies^53^, suggesting this gene is important in conferring tolerance to high concentrations of zinc.

**Figure 7.**
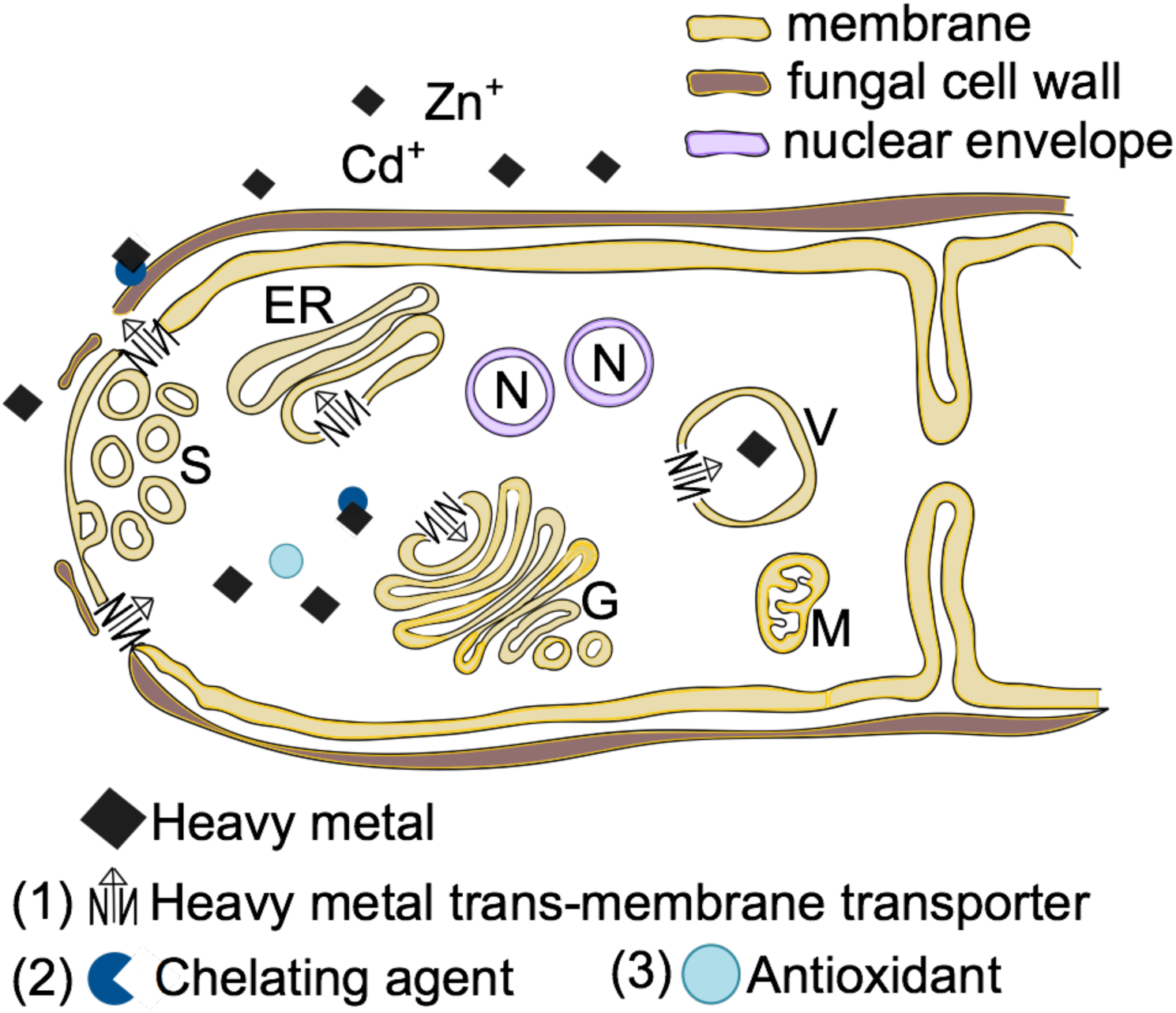
Model of heavy metal tolerance in *S. luteus*, including metal exclusion, storage, immobilization, and detoxification. (1) Exclusion/storage-Heavy metal ion uptake from soil occurs through transporters located in the plasma membrane that can be regulated to adjust metal influx. Heavy metal toxicity induces the release of chelating agents to the outside the cell wall that also restrict metal ion intake. (2) Immobilization-Upon entering the cell, metal ions can be chelated in the cytosol, or actively pumped into vesicles and/or the vacuole by transmembrane transporters. Trapped ions are either stored or shuttled in vesicles to the plasma membrane and extruded from the cell. (3) ROS Detoxification-Antioxidant agents relieve oxidative stress caused by reactive oxygen species (ROS) through reducing agents like reductases or membrane proteins such as Cytochromes p450. Table 1 shows the list of candidate genes putatively involved in S. luteus metal tolerance per functional category. (S=Spitzenkoerper; ER=endoplasmic reticulum; G=Golgi apparatus; V=vesicle/fungal vacuole; N=nucleus; M=mitochondrion).

Many candidate genes were related to metal immobilization, including S-adenosyl-L-methionine methyltransferases, which are known components of the nicotianamine synthesis^82^, a prominent chelating agent of cadmium in fungi^42,82^. Candidate genes also fell in the metal detoxification category, a general and necessary component of cell stress response^36^. Most the differentiated detoxification candidate genes are known to be involved in general stress responses and not specifically involved in metal homeostasis such as manganese/iron superoxide dismutase and glutathione S-transferase. This result is expected, since cells have overarching responses that protect against oxidative stress caused by many factors, including exposure to heavy metals^79^. Indeed, fifteen of our candidate genes under selection were annotated as Cytochromes p450, a numerous family of enzymes that are present in all kingdoms. These membrane proteins perform many functions, including acting as antioxidants and conferring tolerance to pollutants in many organisms^63,84^.

In conclusion, our results show that *S. luteus* is undergoing incipient local adaptation to heavy metal pollution, governed by many genetic variants of small effects. Variants stem from standing genetic variation, with both allelic and copy-number polymorphisms conferring tolerance to soil heavy metal pollution. We also found that *S. luteus* heavy metal tolerance relies on multiple strategies for metal homeostasis (exclusion, storage, immobilization, and ROS detoxification) that involve a combination of genes and pathways. Candidate genes involved in metal transmembrane transport were the main targets of heavy metal selection and further research is needed to confirm the role of all candidate genes in metal tolerance. Given the mycorrhizal habit of *S. luteus*, future investigations should also focus on the interaction between fungal metal tolerance and its pine partners to provide a better understanding of how environmental adaptation occurs in symbiotic organisms.

## Methods

### Culture collection, DNA extraction, and sequencing

*Suillus luteus* cultures were obtained from fruitbodies (mushrooms) collected in heavy metal polluted and unpolluted locations in Belgium (Fig. 1, Table S1). Six localities (three polluted and three non-polluted) were visited at different times during sampling campaigns from fall 1998 to fall 2016. The polluted sites are mainly enriched in zinc and cadmium and, to a lesser extent, in copper, lead, and arsenic^54,85^. The source of the pollution are now decommissioned zinc smelters established ~150 years ago^54^. Fruitbodies were collected at least 10m apart to ensure sampling of different genets and isolated in culture. Isolates were maintained at Hasselt University on solid modified Fries medium, ideal for growing mycorrhizal fungi. We selected ^18^ *S. luteus* isolates from unpolluted and 20 from polluted soils, ranging four to eight isolates per sampled site (Fig. 1). We extracted DNA from cultured mycelia of each isolate using a CTAB/phenol/chloroform protocol (modified from Liao, et al. ^86^). Whole genomes were sequenced with Illumina technology on a HiSeq-2500 1TB model, paired-end with a 600-bp insert size, at the Joint Genome Institute and the raw reads are available in the Short Read Archive (Table S5).

### Read filtering, mapping, genome coverage, and variant calling

Sequences were cleaned by trimming low-quality reads and removing contaminants. Raw reads were filtered with BBmap in the BBTools package^87^. We trimmed all known Illumina adapter sequences, discarded reads with more than 0 ambiguous bases (Ns), and shorter than ^41^ bases after trimming. Read pairs were discarded if both were shorter than 0.33 of the original length after trimming. Retained reads were compared to ‘human’, ‘cat’, ‘dog’, ‘mouse’, and ‘microbes’ databases to remove potential contaminations.

We mapped the reads of each individual sample to the *Suillus luteus* reference genome^88^ (size=44.49 Mb), using Bowtie2^89^. We sorted the mapped alignments in Picard by genomic coordinate and used the same program to add read groups based on read and Illumina run information (Table S5), as well as delete PCR duplicates. We calculated average coverage per genome using the depth of coverage tool in GATK^90^. We called variants in GATK using Haplotype caller in gvcf mode. All samples were genotyped together and single nucleotide polymorphisms (SNPs) were called using the GenotypeGVCF GATK tool. We removed non-biallelic SNPs in GATK, as well as all SNPs with more than 200x or less than 5x coverage with minimum quality above 30 using VCFTools^91^. Using the egglib^92^ package in Python we took the high quality SNPs and the reference genome and created fasta files with in-house scripts (available at: https://github.com/abazzical/sluteus.2019) for each sample.

### Population structure and summary statistics

We investigated the existence of population structure using fastSTRUCTURE^93^ (specifying K as ‘1’, ‘2’, ‘3’, ‘4’, and ‘38’), and by performing a principal components analysis implemented in the R package SNPrelate^94^. We calculated summary statistics on the whole population and groups of isolates from polluted and unpolluted locations. We calculated and Tajima’s D in sci-kit allel^95^ Python package.

### Allelic divergence measures and case/control genome-wide association

To detect genes important for heavy metal tolerance we investigated allelic divergence between isolates from polluted and unpolluted soils. We calculated allele frequency differences (F_ST_) with VCFTools^91^ and the average number of nucleotide differences (d_XY_) using the egglib^92^ package implemented in Python. We estimated both F_ST_ and d_XY_ across 5kb genomic windows. We also implemented a genome-wide association analysis to assess whether specific variants were significantly associated with occurrence in polluted soils. We investigated SNPs associated with contaminated soils by performing a case/control genome-wide association study in PLINK^59^, scoring isolates from polluted soils as ‘case’ and isolates from unpolluted soils as ‘control’. We adjusted the p-values using a Bonferroni correction and a permutation test with 1000 replicates in PLINK^96^.

### Copy-number estimates per isolate and copy-number variation

To detect potential genes involved in heavy metal tolerance based on copy-number variation, we estimated the number of each gene copies for each isolate. We used the software Control-FreeC^97^ to estimate the number of copies based on each isolate’s clean reads. We assumed diploidy of our samples and followed the Steenwyk, et al. ^98^ protocol (see Supplementary Methods and GitHub for detailed settings: https://github.com/abazzical/sluteus.2019). Control-FreeC estimated the number of copies for 250 base pair windows. To identify copy-number variants between isolates in the population from polluted and unpolluted soils, we measured the amount of variance in copies (V_ST_) following Steenwyk, et al. ^98^.

### Gene ontology enrichment analyses

We performed GO enrichment analyses to detect GO terms significantly enriched in the most diverged regions across the *S. luteus* genomes from polluted and non-polluted soils. We used gene ontology (GO) terms based on the annotated reference genome of *Suillus luteus*^88^.We investigated the top 5% differentiated genes for three divergence measures, F_ST_, d_XY_ and copy-number variation and used the Cytoscape^99^ App ClueGO^100^ to describe gene functions. We conducted separate analysis for each divergence measure using default settings and searched for enriched clusters the ‘biological processes’ GO category^101^. We established significance of GO term enrichment by implementing the ClueGO statistics using a two-sided hypergeometric test and Benjamini-Hochberg corrected p-values.

## Acknowledgements

This work was supported by the Joint Genome Institute Community Science Program (award JGI 502931 to NN) and by NSF grant DEB-1554181 (to RV). JVC and JR were funded by the Research Foundation Flanders (FWO project G079213N to JVC) and Vrije Universiteit Brussel (starting grant JR). LC was supported by a Flanders Innovation & Entrepreneurships PhD fellowship (IWT project 141461). SB was supported by the Montana’s Agricultural Experiment Station. NN was supported by Hawaii’s Agricultural Experiment Station. We thank P. Gladieux for assisting in data analyses, G. Bindea for her help in including *Suillus luteus* in ClueGO, and H-L. Liao for helpful comments on the manuscript.

