## Supplementary figures and tables for "Incipient local adaptation in a fungus: evolution of heavy metal tolerance through allelic and copy-number variation"

### Supplementary Information:

#### Supplementary Figures

See GitHub (https://github.com/abazzical/sluteus.2019) for commands used in the analyses.

### Supplementary Figures

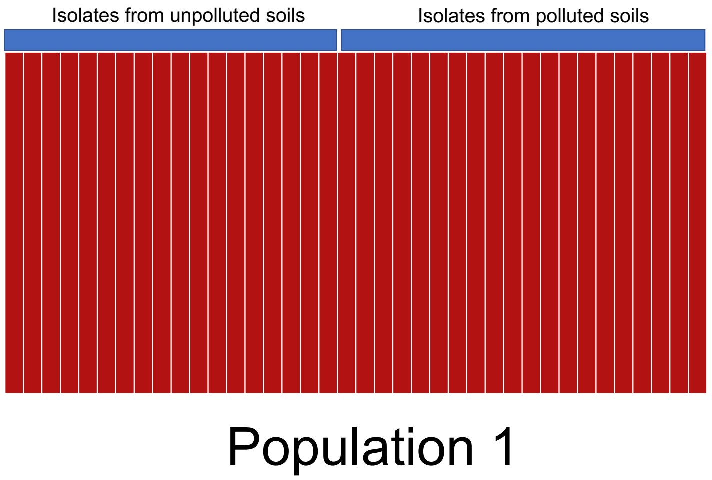

Figure S 1 The STRUCTURE plot showing the absence of population structure.

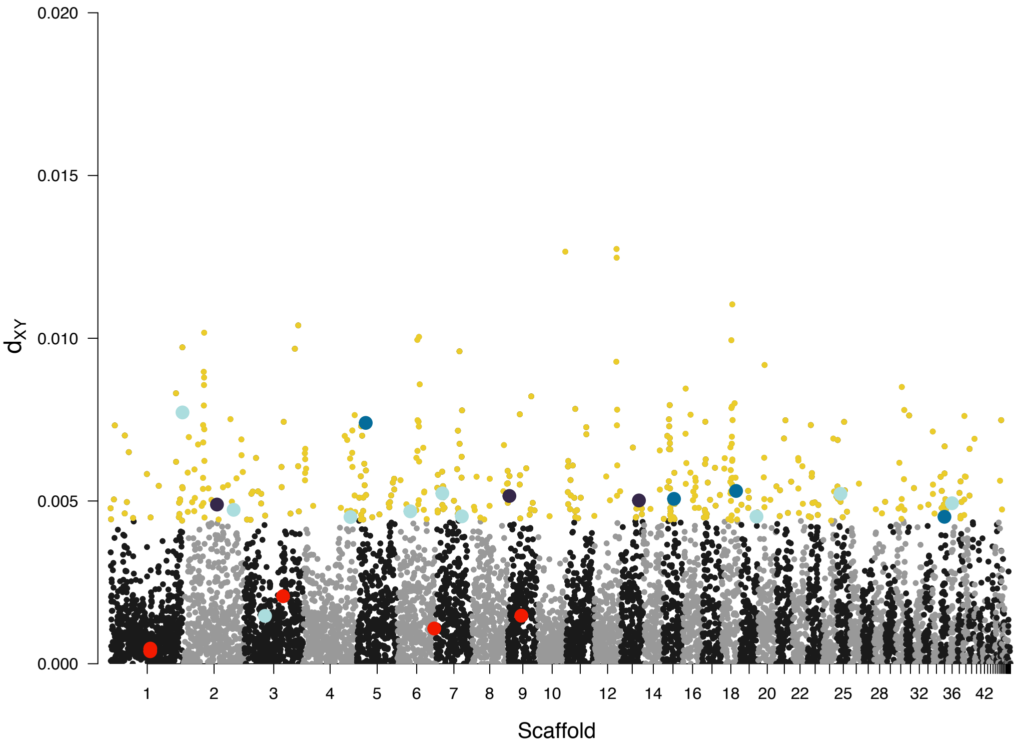

*Figure S 2* The average dxy values across 5kb windows between S. luteus from polluted and non-polluted soils.

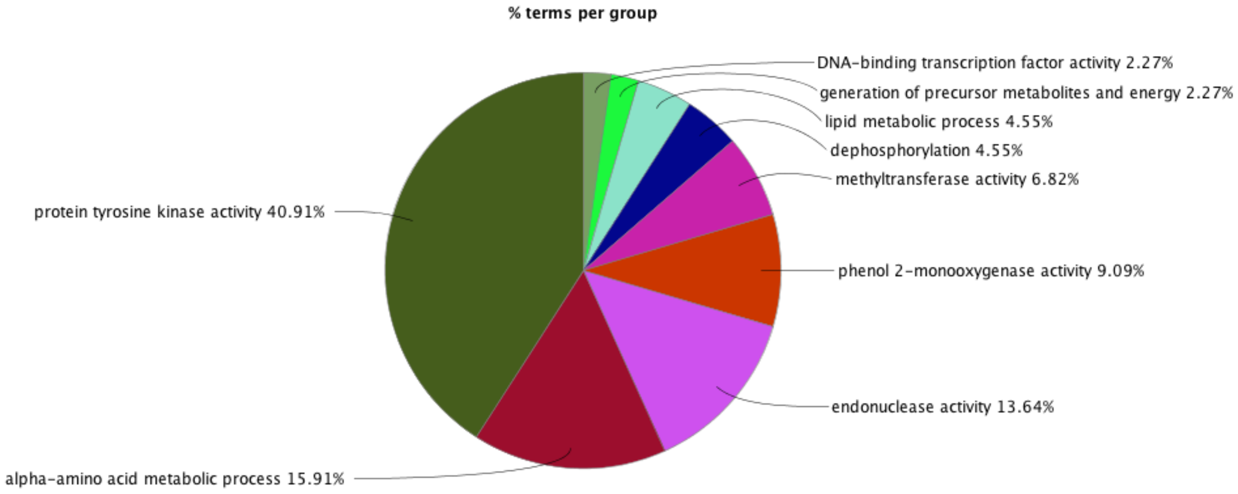

*Figure S 3 Pie chart of the significant gene ontology (GO) enriched terms in the top 5% dxy values, with respective percentages.*

### Supplementary Tables

*Table S 1 Fungal isolates metadata and genome coverage information.*

| Sample name | Collection year | soil Zn | soil Cd | locality | Total no. reads | Overall genome coverage | Total number of reads | % bases above 15x |
|---|---|---|---|---|---|---|---|---|
| UH-Slu-B10 | 2016 | - | - | Bilzen, BE | 2627127607 | 59.05 | 2627127607 | 88.7 |
| UH-Slu-B12 | 2016 | - | - | Bilzen, BE | 2155505492 | 48.45 | 2155505492 | 87.9 |
| UH-Slu-B2a | 2016 | - | - | Bilzen, BE | 2612030695 | 58.72 | 2612030695 | 89.7 |
| UH-Slu-B3c | 2016 | - | - | Bilzen, BE | 2240392646 | 50.36 | 2240392646 | 86.2 |
| UH-Slu-B4a | 2016 | - | - | Bilzen, BE | 2354867057 | 52.93 | 2354867057 | 88.1 |
| UH-Slu-B6 | 2016 | - | - | Bilzen, BE | 2376547549 | 53.42 | 2376547549 | 89.2 |
| UH-Slu-B7b | 2016 | - | - | Bilzen, BE | 2427660950 | 54.57 | 2427660950 | 89 |
| UH-Slu-B9a | 2016 | - | - | Bilzen, BE | 2211966292 | 49.72 | 2211966292 | 88.6 |
| UH-Slu-DS16 | 2003 | + | + | Dilsen, BE | 1097924730 | 24.68 | 1097924730 | 77.9 |
| UH-Slu-DS3 | 2003 | + | + | Dilsen, BE | 721145426 | 16.21 | 721145426 | 55.7 |
| UH-Slu-DS4 | 2003 | + | + | Dilsen, BE | 1450668437 | 32.61 | 1450668437 | 83.3 |
| UH-Slu-DS5 | 2003 | + | + | Dilsen, BE | 1688292704 | 37.95 | 1688292704 | 84.9 |
| UH-Slu-DS7 | 2003 | + | + | Dilsen, BE | 1140880168 | 25.65 | 1140880168 | 77.9 |
| UH-Slu-DS9 | 2003 | + | + | Dilsen, BE | 1693289518 | 38.06 | 1693289518 | 84 |
| UH-Slu-Ew1 | 2001 | - | - | Eksel, BE | 1529627052 | 34.38 | 1529627052 | 84 |
| UH-Slu-Ew2 | 2001 | - | - | Eksel, BE | 990223356 | 22.26 | 990223356 | 73.5 |
| UH-Slu-Ew3 | 2001 | - | - | Eksel, BE | 7233239115 | 162.59 | 7233239115 | 92.7 |
| UH-Slu-Ew7 | 2001 | - | - | Eksel, BE | 1331854648 | 29.94 | 1331854648 | 81.7 |
| UH-Slu-LM12 | 2000 | + | + | Lommel, BE | 1126641088 | 25.33 | 1126641088 | 76.9 |
| UH-Slu-LM3 | 2000 | + | + | Lommel, BE | 1095475838 | 24.62 | 1095475838 | 76.6 |
| UH-Slu-LM5 | 2000 | + | + | Lommel, BE | 1003925030 | 22.57 | 1003925030 | 73.7 |
| UH-Slu-LmD7 | 2011 | + | + | Lommel, BE | 1382140865 | 31.07 | 1382140865 | 82.3 |
| UH-Slu-Sl1 | 1995 | + | + | Lommel, BE | 908788591 | 20.43 | 908788591 | 71.6 |
| UH-Slu-Sl2 | 1995 | + | + | Lommel, BE | 638673094 | 14.36 | 638673094 | 44.2 |
| UH-Slu-Sl20 | 1998 | + | + | Lommel, BE | 842990653 | 18.95 | 842990653 | 66.7 |
| UH-Slu-Sl22 | 1998 | + | + | Lommel, BE | 920336661 | 20.69 | 920336661 | 71.3 |
| UH-Slu-Sl24 | 1998 | + | + | Lommel, BE | 958302965 | 21.54 | 958302965 | 73.5 |
| UH-Slu-N3 | 2000 | + | + | Neerpelt, BE | 810255631 | 18.21 | 810255631 | 64.1 |
| UH-Slu-N4 | 2000 | + | + | Neerpelt, BE | 723116214 | 16.25 | 723116214 | 56.3 |
| UH-Slu-N5 | 2000 | + | + | Neerpelt, BE | 885325848 | 19.9 | 885325848 | 70.3 |
| UH-Slu-Na2 | 2000 | + | + | Neerpelt, BE | 1327518074 | 29.84 | 1327518074 | 80.8 |

| | | | | | | | | |
|---|---|---|---|---|---|---|---|---|
| UH-Slu-Na5 | 2000 | + | + | Neerpelt, BE | 1107682223 | 24.9 | 1107682223 | 76.2 |
| UH-Slu-P2 | 2000 | - | - | Paal, BE | 991366408 | 22.28 | 991366408 | 74.2 |
| UH-Slu-P8 | 2000 | - | - | Paal, BE | 1225673854 | 27.55 | 1225673854 | 78.2 |
| UH-Slu-PD13 | 2011 | - | - | Paal, BE | 1095944827 | 24.64 | 1095944827 | 77.6 |
| UH-Slu-PD14 | 2011 | - | - | Paal, BE | 966698558 | 21.73 | 966698558 | 73.7 |
| UH-Slu-Sl31 | 1998 | - | - | Paal, BE | 825274367 | 18.55 | 825274367 | 64.7 |
| UH-Slu-Sl33 | 1998 | - | - | Paal, BE | 604532312 | 13.59 | 604532312 | 40.4 |

*Table S 2 Significant GOterms ‘biological process’ for top 5% regions of FST*

| GOID | GOTerm | Ontology Source | Term PValue | Term PValue Corrected with Bonferroni step down | Group PValue | Group PValue Corrected with Bonferroni step down | GOLevels | GOGroups | % Associated Genes | Nr. Genes |
|---|---|---|---|---|---|---|---|---|---|---|
| GO:0004527 | exonuclease activity | GO_BiologicalProcess-Custom-GOA_05.03.2019_00h00 | 0.07 | 1.00 | 0.07 | 0.99 | [7, 8] | Group00 | 11.76 | 4.00 |
| GO:0022607 | cellular component assembly | GO_BiologicalProcess-Custom-GOA_05.03.2019_00h00 | 0.49 | 1.00 | 0.49 | 1.00 | [3] | Group01 | 6.00 | 3.00 |
| GO:0044255 | cellular lipid metabolic process | GO_BiologicalProcess-Custom-GOA_05.03.2019_00h00 | 1.00 | 1.00 | 1.00 | 1.00 | [3, 4] | Group02 | 4.30 | 4.00 |
| GO:0044262 | cellular carbohydrate metabolic process | GO_BiologicalProcess-Custom-GOA_05.03.2019_00h00 | 0.08 | 1.00 | 0.08 | 1.00 | [3, 4] | Group03 | 10.81 | 4.00 |
| GO:0051186 | cofactor metabolic process | GO_BiologicalProcess-Custom-GOA_05.03.2019_00h00 | 1.00 | 1.00 | 1.00 | 1.00 | [3] | Group04 | 4.35 | 3.00 |
| GO:0005975 | carbohydrate metabolic process | GO_BiologicalProcess-Custom-GOA_05.03.2019_00h00 | 0.03 | 1.00 | 0.03 | 0.43 | [3] | Group05 | 8.29 | 15.00 |
| GO:0004497 | monooxygenase activity | GO_BiologicalProcess-Custom-GOA_05.03.2019_00h00 | 0.85 | 1.00 | 0.85 | 1.00 | [4] | Group06 | 4.71 | 8.00 |
| GO:0009132 | nucleoside diphosphate metabolic process | GO_BiologicalProcess-Custom-GOA_05.03.2019_00h00 | 0.07 | 1.00 | 0.07 | 0.94 | [5, 6, 7] | Group07 | 14.29 | 3.00 |
| GO:0016651 | oxidoreductase activity, acting on NAD(P)H | GO_BiologicalProcess-Custom-GOA_05.03.2019_00h00 | 0.74 | 1.00 | 0.74 | 1.00 | [4] | Group08 | 5.45 | 3.00 |
| GO:0017144 | drug metabolic process | GO_BiologicalProcess-Custom-GOA_05.03.2019_00h00 | 1.00 | 1.00 | 0.84 | 1.00 | [3] | Group09 | 4.35 | 4.00 |
| GO:1901137 | carbohydrate derivative biosynthetic process | GO_BiologicalProcess-Custom-GOA_05.03.2019_00h00 | 1.00 | 1.00 | 0.84 | 1.00 | [4] | Group09 | 4.44 | 4.00 |

| | | | | | | | | | | |
|---|---|---|---|---|---|---|---|---|---|---|
| GO:0005996 | monosaccharide metabolic process | GO_BiologicalProcess-Custom-GOA_05.03.2019_00h00 | 0.03 | 1.00 | 0.03 | 0.53 | [3, 4] | Group10 | 18.75 | 3.00 |
| GO:0019318 | hexose metabolic process | GO_BiologicalProcess-Custom-GOA_05.03.2019_00h00 | 0.03 | 1.00 | 0.03 | 0.53 | [4, 5] | Group10 | 20.00 | 3.00 |
| GO:0034248 | regulation of cellular amide metabolic process | GO_BiologicalProcess-Custom-GOA_05.03.2019_00h00 | 0.73 | 1.00 | 0.74 | 1.00 | [4, 5] | Group11 | 5.66 | 3.00 |
| GO:0010608 | posttranscriptional regulation of gene expression | GO_BiologicalProcess-Custom-GOA_05.03.2019_00h00 | 0.74 | 1.00 | 0.74 | 1.00 | [5, 6] | Group11 | 5.56 | 3.00 |
| GO:0006413 | translational initiation | GO_BiologicalProcess-Custom-GOA_05.03.2019_00h00 | 0.23 | 1.00 | 0.74 | 1.00 | [3, 6, 7, 8] | Group11 | 8.11 | 3.00 |
| GO:0006417 | regulation of translation | GO_BiologicalProcess-Custom-GOA_05.03.2019_00h00 | 0.73 | 1.00 | 0.74 | 1.00 | [5, 6, 7, 8] | Group11 | 5.66 | 3.00 |
| GO:0008168 | methyltransferase activity | GO_BiologicalProcess-Custom-GOA_05.03.2019_00h00 | 0.82 | 1.00 | 1.00 | 1.00 | [3] | Group12 | 4.67 | 5.00 |
| GO:0022613 | ribonucleoprotein complex biogenesis | GO_BiologicalProcess-Custom-GOA_05.03.2019_00h00 | 0.49 | 1.00 | 1.00 | 1.00 | [3] | Group12 | 6.12 | 3.00 |
| GO:0008170 | N-methyltransferase activity | GO_BiologicalProcess-Custom-GOA_05.03.2019_00h00 | 0.44 | 1.00 | 1.00 | 1.00 | [4] | Group12 | 7.14 | 3.00 |
| GO:0042254 | ribosome biogenesis | GO_BiologicalProcess-Custom-GOA_05.03.2019_00h00 | 0.48 | 1.00 | 1.00 | 1.00 | [4] | Group12 | 6.25 | 3.00 |
| GO:0016705 | oxidoreductase activity, acting on paired donors, with incorporation or reduction of molecular oxygen | GO_BiologicalProcess-Custom-GOA_05.03.2019_00h00 | 1.00 | 1.00 | 1.00 | 1.00 | [4] | Group13 | 4.08 | 4.00 |
| GO:0051213 | dioxygenase activity | GO_BiologicalProcess-Custom-GOA_05.03.2019_00h00 | 0.26 | 1.00 | 1.00 | 1.00 | [4] | Group13 | 7.69 | 3.00 |
| GO:0016706 | 2-oxoglutarate-dependent dioxygenase activity | GO_BiologicalProcess-Custom-GOA_05.03.2019_00h00 | 0.16 | 1.00 | 1.00 | 1.00 | [5] | Group13 | 9.68 | 3.00 |
| GO:0018208 | peptidyl-proline modification | GO_BiologicalProcess-Custom-GOA_05.03.2019_00h00 | 0.26 | 1.00 | 1.00 | 1.00 | [7, 8] | Group13 | 7.69 | 3.00 |
| GO:0023051 | regulation of signaling | GO_BiologicalProcess-Custom-GOA_05.03.2019_00h00 | 1.00 | 1.00 | 0.78 | 1.00 | [2, 3] | Group14 | 4.23 | 3.00 |
| GO:0048583 | regulation of response to stimulus | GO_BiologicalProcess-Custom-GOA_05.03.2019_00h00 | 1.00 | 1.00 | 0.78 | 1.00 | [2, 3] | Group14 | 4.23 | 3.00 |

| | | | | | | | | | | |
|---|---|---|---|---|---|---|---|---|---|---|
| GO:0010646 | regulation of cell communication | GO_BiologicalProcess-Custom-GOA_05.03.2019_00h00 | 1.00 | 1.00 | 0.78 | 1.00 | [3, 4] | Group14 | 4.23 | 3.00 |
| GO:0005085 | guanyl-nucleotide exchange factor activity | GO_BiologicalProcess-Custom-GOA_05.03.2019_00h00 | 0.46 | 1.00 | 0.78 | 1.00 | [4] | Group14 | 6.67 | 3.00 |
| GO:0009966 | regulation of signal transduction | GO_BiologicalProcess-Custom-GOA_05.03.2019_00h00 | 1.00 | 1.00 | 0.78 | 1.00 | [3, 4, 5] | Group14 | 4.23 | 3.00 |
| GO:1902531 | regulation of intracellular signal transduction | GO_BiologicalProcess-Custom-GOA_05.03.2019_00h00 | 1.00 | 1.00 | 0.78 | 1.00 | [4, 5, 6] | Group14 | 4.48 | 3.00 |
| GO:0006508 | proteolysis | GO_BiologicalProcess-Custom-GOA_05.03.2019_00h00 | 0.07 | 1.00 | 0.07 | 0.94 | [4, 5] | Group15 | 6.96 | 16.00 |
| GO:0008233 | peptidase activity | GO_BiologicalProcess-Custom-GOA_05.03.2019_00h00 | 0.10 | 1.00 | 0.07 | 0.94 | [5, 6] | Group15 | 7.18 | 13.00 |
| GO:0070011 | peptidase activity, acting on L-amino acid peptides | GO_BiologicalProcess-Custom-GOA_05.03.2019_00h00 | 0.25 | 1.00 | 0.07 | 0.94 | [6, 7] | Group15 | 6.43 | 11.00 |
| GO:0004175 | endopeptidase activity | GO_BiologicalProcess-Custom-GOA_05.03.2019_00h00 | 0.55 | 1.00 | 0.07 | 0.94 | [7, 8] | Group15 | 5.97 | 4.00 |
| GO:0008234 | cysteine-type peptidase activity | GO_BiologicalProcess-Custom-GOA_05.03.2019_00h00 | 0.11 | 1.00 | 0.07 | 0.94 | [7, 8] | Group15 | 10.00 | 4.00 |
| GO:0008238 | exopeptidase activity | GO_BiologicalProcess-Custom-GOA_05.03.2019_00h00 | 0.18 | 1.00 | 0.07 | 0.94 | [7, 8] | Group15 | 9.38 | 3.00 |
| GO:0070001 | aspartic-type peptidase activity | GO_BiologicalProcess-Custom-GOA_05.03.2019_00h00 | 0.15 | 1.00 | 0.07 | 0.94 | [7, 8] | Group15 | 10.00 | 3.00 |
| GO:0004190 | aspartic-type endopeptidase activity | GO_BiologicalProcess-Custom-GOA_05.03.2019_00h00 | 0.15 | 1.00 | 0.07 | 0.94 | [8, 9] | Group15 | 10.00 | 3.00 |
| GO:0033554 | cellular response to stress | GO_BiologicalProcess-Custom-GOA_05.03.2019_00h00 | 0.62 | 1.00 | 0.92 | 1.00 | [3] | Group16 | 5.32 | 5.00 |
| GO:0006974 | cellular response to DNA damage stimulus | GO_BiologicalProcess-Custom-GOA_05.03.2019_00h00 | 1.00 | 1.00 | 0.92 | 1.00 | [4] | Group16 | 4.17 | 3.00 |
| GO:0090304 | nucleic acid metabolic process | GO_BiologicalProcess-Custom-GOA_05.03.2019_00h00 | 1.00 | 1.00 | 0.92 | 1.00 | [4, 5] | Group16 | 4.41 | 27.00 |
| GO:0006259 | DNA metabolic process | GO_BiologicalProcess-Custom-GOA_05.03.2019_00h00 | 1.00 | 1.00 | 0.92 | 1.00 | [4, 5, 6] | Group16 | 4.57 | 8.00 |
| GO:0090305 | nucleic acid phosphodiester bond hydrolysis | GO_BiologicalProcess-Custom-GOA_05.03.2019_00h00 | 0.00 | 0.00 | 0.92 | 1.00 | [5, 6] | Group16 | 12.84 | 19.00 |
| GO:0006281 | DNA repair | GO_BiologicalProcess-Custom- | 1.00 | 1.00 | 0.92 | 1.00 | [5, 6, 7] | Group16 | 4.23 | 3.00 |

| | | | | | | | | | | |
|---|---|---|---|---|---|---|---|---|---|---|
| | | GOA_05.03.2019_00h00 | | | | | | | | |
| GO:0004518 | nuclease activity | GO_BiologicalProcess-Custom-GOA_05.03.2019_00h00 | 0.00 | 0.00 | 0.92 | 1.00 | [6, 7] | Group16 | 12.84 | 19.00 |
| GO:0004536 | deoxyribonuclease activity | GO_BiologicalProcess-Custom-GOA_05.03.2019_00h00 | 0.43 | 1.00 | 0.92 | 1.00 | [5, 6, 7, 8] | Group16 | 7.32 | 3.00 |
| GO:0044248 | cellular catabolic process | GO_BiologicalProcess-Custom-GOA_05.03.2019_00h00 | 0.60 | 1.00 | 0.81 | 1.00 | [3] | Group17 | 5.62 | 5.00 |
| GO:1901575 | organic substance catabolic process | GO_BiologicalProcess-Custom-GOA_05.03.2019_00h00 | 0.81 | 1.00 | 0.81 | 1.00 | [3] | Group17 | 5.00 | 5.00 |
| GO:0009057 | macromolecule catabolic process | GO_BiologicalProcess-Custom-GOA_05.03.2019_00h00 | 0.54 | 1.00 | 0.81 | 1.00 | [4] | Group17 | 6.15 | 4.00 |
| GO:1901565 | organonitrogen compound catabolic process | GO_BiologicalProcess-Custom-GOA_05.03.2019_00h00 | 0.57 | 1.00 | 0.81 | 1.00 | [4] | Group17 | 5.56 | 4.00 |
| GO:0030163 | protein catabolic process | GO_BiologicalProcess-Custom-GOA_05.03.2019_00h00 | 0.29 | 1.00 | 0.81 | 1.00 | [4, 5] | Group17 | 7.84 | 4.00 |
| GO:0044265 | cellular macromolecule catabolic process | GO_BiologicalProcess-Custom-GOA_05.03.2019_00h00 | 0.16 | 1.00 | 0.81 | 1.00 | [4, 5] | Group17 | 8.51 | 4.00 |
| GO:0043632 | modification-dependent macromolecule catabolic process | GO_BiologicalProcess-Custom-GOA_05.03.2019_00h00 | 0.11 | 1.00 | 0.81 | 1.00 | [5, 6] | Group17 | 9.76 | 4.00 |
| GO:0044257 | cellular protein catabolic process | GO_BiologicalProcess-Custom-GOA_05.03.2019_00h00 | 0.11 | 1.00 | 0.81 | 1.00 | [5, 6] | Group17 | 9.76 | 4.00 |
| GO:0051603 | proteolysis involved in cellular protein catabolic process | GO_BiologicalProcess-Custom-GOA_05.03.2019_00h00 | 0.11 | 1.00 | 0.81 | 1.00 | [5, 6, 7] | Group17 | 9.76 | 4.00 |
| GO:0019941 | modification-dependent protein catabolic process | GO_BiologicalProcess-Custom-GOA_05.03.2019_00h00 | 0.11 | 1.00 | 0.81 | 1.00 | [6, 7, 8] | Group17 | 9.76 | 4.00 |
| GO:0006511 | ubiquitin-dependent protein catabolic process | GO_BiologicalProcess-Custom-GOA_05.03.2019_00h00 | 0.11 | 1.00 | 0.81 | 1.00 | [7, 8, 9] | Group17 | 9.76 | 4.00 |
| GO:0006725 | cellular aromatic compound metabolic process | GO_BiologicalProcess-Custom-GOA_05.03.2019_00h00 | 0.51 | 1.00 | 0.23 | 1.00 | [3] | Group18 | 4.05 | 34.00 |
| GO:0034641 | cellular nitrogen compound metabolic process | GO_BiologicalProcess-Custom-GOA_05.03.2019_00h00 | 0.43 | 1.00 | 0.23 | 1.00 | [3] | Group18 | 4.04 | 40.00 |

| | | | | | | | | | | |
|---|---|---|---|---|---|---|---|---|---|---|
| GO:0046483 | heterocycle metabolic process | GO_BiologicalProcess-Custom-GOA_05.03.2019_00h00 | 0.70 | 1.00 | 0.23 | 1.00 | [3] | Group18 | 4.20 | 34.00 |
| GO:1901360 | organic cyclic compound metabolic process | GO_BiologicalProcess-Custom-GOA_05.03.2019_00h00 | 0.51 | 1.00 | 0.23 | 1.00 | [3] | Group18 | 4.08 | 34.00 |
| GO:0006139 | nucleobase-containing compound metabolic process | GO_BiologicalProcess-Custom-GOA_05.03.2019_00h00 | 0.77 | 1.00 | 0.23 | 1.00 | [3, 4] | Group18 | 4.25 | 32.00 |
| GO:0090304 | nucleic acid metabolic process | GO_BiologicalProcess-Custom-GOA_05.03.2019_00h00 | 1.00 | 1.00 | 0.23 | 1.00 | [4, 5] | Group18 | 4.41 | 27.00 |
| GO:0006259 | DNA metabolic process | GO_BiologicalProcess-Custom-GOA_05.03.2019_00h00 | 1.00 | 1.00 | 0.23 | 1.00 | [4, 5, 6] | Group18 | 4.57 | 8.00 |
| GO:0016070 | RNA metabolic process | GO_BiologicalProcess-Custom-GOA_05.03.2019_00h00 | 0.81 | 1.00 | 0.23 | 1.00 | [5, 6] | Group18 | 4.20 | 19.00 |
| GO:0090305 | nucleic acid phosphodiester bond hydrolysis | GO_BiologicalProcess-Custom-GOA_05.03.2019_00h00 | 0.00 | 0.00 | 0.23 | 1.00 | [5, 6] | Group18 | 12.84 | 19.00 |
| GO:0004518 | nuclease activity | GO_BiologicalProcess-Custom-GOA_05.03.2019_00h00 | 0.00 | 0.00 | 0.23 | 1.00 | [6, 7] | Group18 | 12.84 | 19.00 |
| GO:0090501 | RNA phosphodiester bond hydrolysis | GO_BiologicalProcess-Custom-GOA_05.03.2019_00h00 | 0.00 | 0.02 | 0.23 | 1.00 | [6, 7] | Group18 | 14.29 | 12.00 |
| GO:0004519 | endonuclease activity | GO_BiologicalProcess-Custom-GOA_05.03.2019_00h00 | 0.00 | 0.01 | 0.23 | 1.00 | [7, 8] | Group18 | 13.27 | 15.00 |
| GO:0004540 | ribonuclease activity | GO_BiologicalProcess-Custom-GOA_05.03.2019_00h00 | 0.00 | 0.02 | 0.23 | 1.00 | [7, 8] | Group18 | 14.29 | 12.00 |
| GO:0090502 | RNA phosphodiester bond hydrolysis, endonucleolytic | GO_BiologicalProcess-Custom-GOA_05.03.2019_00h00 | 0.00 | 0.02 | 0.23 | 1.00 | [7, 8] | Group18 | 15.94 | 11.00 |
| GO:0004521 | endoribonuclease activity | GO_BiologicalProcess-Custom-GOA_05.03.2019_00h00 | 0.00 | 0.02 | 0.23 | 1.00 | [8, 9] | Group18 | 15.94 | 11.00 |
| GO:0016893 | endonuclease activity, active with either ribo- or deoxyribonucleic acids and producing 5'-phosphomonoesters | GO_BiologicalProcess-Custom-GOA_05.03.2019_00h00 | 0.00 | 0.01 | 0.23 | 1.00 | [8, 9] | Group18 | 16.42 | 11.00 |
| GO:0006793 | phosphorus metabolic process | GO_BiologicalProcess-Custom-GOA_05.03.2019_00h00 | 0.07 | 1.00 | 0.64 | 1.00 | [3] | Group19 | 5.98 | 36.00 |

| | | | | | | | | | | |
|---|---|---|---|---|---|---|---|---|---|---|
| GO:0043170 | macromolecule metabolic process | GO_BiologicalProcess-Custom-GOA_05.03.2019_00h00 | 0.27 | 1.00 | 0.64 | 1.00 | [3] | Group19 | 4.98 | 82.00 |
| GO:1901564 | organonitrogen compound metabolic process | GO_BiologicalProcess-Custom-GOA_05.03.2019_00h00 | 0.69 | 1.00 | 0.64 | 1.00 | [3] | Group19 | 4.33 | 58.00 |
| GO:0019538 | protein metabolic process | GO_BiologicalProcess-Custom-GOA_05.03.2019_00h00 | 0.61 | 1.00 | 0.64 | 1.00 | [3, 4] | Group19 | 4.82 | 52.00 |
| GO:0044260 | cellular macromolecule metabolic process | GO_BiologicalProcess-Custom-GOA_05.03.2019_00h00 | 0.74 | 1.00 | 0.64 | 1.00 | [3, 4] | Group19 | 4.34 | 53.00 |
| GO:0006796 | phosphate-containing compound metabolic process | GO_BiologicalProcess-Custom-GOA_05.03.2019_00h00 | 0.08 | 1.00 | 0.64 | 1.00 | [4] | Group19 | 5.94 | 35.00 |
| GO:0043412 | macromolecule modification | GO_BiologicalProcess-Custom-GOA_05.03.2019_00h00 | 0.54 | 1.00 | 0.64 | 1.00 | [4] | Group19 | 4.99 | 33.00 |
| GO:0036211 | protein modification process | GO_BiologicalProcess-Custom-GOA_05.03.2019_00h00 | 0.47 | 1.00 | 0.64 | 1.00 | [4, 5] | Group19 | 5.06 | 32.00 |
| GO:0044267 | cellular protein metabolic process | GO_BiologicalProcess-Custom-GOA_05.03.2019_00h00 | 1.00 | 1.00 | 0.64 | 1.00 | [4, 5] | Group19 | 4.53 | 40.00 |
| GO:0016310 | phosphorylation | GO_BiologicalProcess-Custom-GOA_05.03.2019_00h00 | 0.00 | 0.26 | 0.64 | 1.00 | [5] | Group19 | 7.63 | 29.00 |
| GO:0006464 | cellular protein modification process | GO_BiologicalProcess-Custom-GOA_05.03.2019_00h00 | 0.47 | 1.00 | 0.64 | 1.00 | [5, 6] | Group19 | 5.06 | 32.00 |
| GO:0016301 | kinase activity | GO_BiologicalProcess-Custom-GOA_05.03.2019_00h00 | 0.01 | 0.32 | 0.64 | 1.00 | [6] | Group19 | 7.67 | 28.00 |
| GO:0006468 | protein phosphorylation | GO_BiologicalProcess-Custom-GOA_05.03.2019_00h00 | 0.00 | 0.15 | 0.64 | 1.00 | [6, 7] | Group19 | 8.33 | 25.00 |
| GO:0018193 | peptidyl-amino acid modification | GO_BiologicalProcess-Custom-GOA_05.03.2019_00h00 | 0.02 | 1.00 | 0.64 | 1.00 | [6, 7] | Group19 | 8.54 | 14.00 |
| GO:0004672 | protein kinase activity | GO_BiologicalProcess-Custom-GOA_05.03.2019_00h00 | 0.00 | 0.15 | 0.64 | 1.00 | [7, 8] | Group19 | 8.33 | 25.00 |
| GO:0018212 | peptidyl-tyrosine modification | GO_BiologicalProcess-Custom-GOA_05.03.2019_00h00 | 0.00 | 0.20 | 0.64 | 1.00 | [7, 8] | Group19 | 12.50 | 10.00 |
| GO:0018108 | peptidyl-tyrosine phosphorylation | GO_BiologicalProcess-Custom-GOA_05.03.2019_00h00 | 0.00 | 0.20 | 0.64 | 1.00 | [7, 8, 9] | Group19 | 12.50 | 10.00 |
| GO:0004674 | protein serine/threonine kinase activity | GO_BiologicalProcess-Custom-GOA_05.03.2019_00h00 | 0.03 | 1.00 | 0.64 | 1.00 | [8, 9] | Group19 | 7.77 | 15.00 |

| | | | | | | | | | | |
|---|---|---|---|---|---|---|---|---|---|---|
| GO:0004713 | protein tyrosine kinase activity | GO_BiologicalProcess-Custom-GOA_05.03.2019_00h00 | 0.00 | 0.20 | 0.64 | 1.00 | [8, 9, 10] | Group19 | 12.50 | 10.00 |
| GO:0006810 | transport | GO_BiologicalProcess-Custom-GOA_05.03.2019_00h00 | 0.73 | 1.00 | 0.54 | 1.00 | [3] | Group20 | 4.13 | 21.00 |
| GO:0006996 | organelle organization | GO_BiologicalProcess-Custom-GOA_05.03.2019_00h00 | 1.00 | 1.00 | 0.54 | 1.00 | [3] | Group20 | 4.38 | 7.00 |
| GO:0008104 | protein localization | GO_BiologicalProcess-Custom-GOA_05.03.2019_00h00 | 0.84 | 1.00 | 0.54 | 1.00 | [3] | Group20 | 4.64 | 7.00 |
| GO:0045184 | establishment of protein localization | GO_BiologicalProcess-Custom-GOA_05.03.2019_00h00 | 0.84 | 1.00 | 0.54 | 1.00 | [3, 4] | Group20 | 4.70 | 7.00 |
| GO:0016192 | vesicle-mediated transport | GO_BiologicalProcess-Custom-GOA_05.03.2019_00h00 | 0.25 | 1.00 | 0.54 | 1.00 | [4] | Group20 | 7.14 | 5.00 |
| GO:0051276 | chromosome organization | GO_BiologicalProcess-Custom-GOA_05.03.2019_00h00 | 1.00 | 1.00 | 0.54 | 1.00 | [4] | Group20 | 4.20 | 5.00 |
| GO:0055085 | transmembrane transport | GO_BiologicalProcess-Custom-GOA_05.03.2019_00h00 | 0.65 | 1.00 | 0.54 | 1.00 | [4] | Group20 | 4.98 | 13.00 |
| GO:0071702 | organic substance transport | GO_BiologicalProcess-Custom-GOA_05.03.2019_00h00 | 1.00 | 1.00 | 0.54 | 1.00 | [4] | Group20 | 4.30 | 11.00 |
| GO:0071705 | nitrogen compound transport | GO_BiologicalProcess-Custom-GOA_05.03.2019_00h00 | 0.73 | 1.00 | 0.54 | 1.00 | [4] | Group20 | 4.85 | 10.00 |
| GO:0022857 | transmembrane transporter activity | GO_BiologicalProcess-Custom-GOA_05.03.2019_00h00 | 0.64 | 1.00 | 0.54 | 1.00 | [5] | Group20 | 5.00 | 13.00 |
| GO:0042886 | amide transport | GO_BiologicalProcess-Custom-GOA_05.03.2019_00h00 | 0.84 | 1.00 | 0.54 | 1.00 | [5] | Group20 | 4.70 | 7.00 |
| GO:0071103 | DNA conformation change | GO_BiologicalProcess-Custom-GOA_05.03.2019_00h00 | 0.76 | 1.00 | 0.54 | 1.00 | [5] | Group20 | 4.84 | 3.00 |
| GO:0090662 | ATP hydrolysis coupled transmembrane transport | GO_BiologicalProcess-Custom-GOA_05.03.2019_00h00 | 0.76 | 1.00 | 0.54 | 1.00 | [5] | Group20 | 4.84 | 3.00 |
| GO:0015031 | protein transport | GO_BiologicalProcess-Custom-GOA_05.03.2019_00h00 | 0.84 | 1.00 | 0.54 | 1.00 | [4, 5, 6, 7] | Group20 | 4.76 | 7.00 |
| GO:0015833 | peptide transport | GO_BiologicalProcess-Custom-GOA_05.03.2019_00h00 | 0.84 | 1.00 | 0.54 | 1.00 | [5, 6] | Group20 | 4.70 | 7.00 |
| GO:0022804 | active transmembrane transporter activity | GO_BiologicalProcess-Custom-GOA_05.03.2019_00h00 | 1.00 | 1.00 | 0.54 | 1.00 | [6] | Group20 | 4.19 | 7.00 |

| | | | | | | | | | | |
|---|---|---|---|---|---|---|---|---|---|---|
| GO:0008565 | protein transporter activity | GO_BiologicalProcess-Custom-GOA_05.03.2019_00h00 | 0.47 | 1.00 | 0.54 | 1.00 | [5, 6, 7, 8] | Group20 | 6.38 | 3.00 |
| GO:0099131 | ATP hydrolysis coupled ion transmembrane transport | GO_BiologicalProcess-Custom-GOA_05.03.2019_00h00 | 0.76 | 1.00 | 0.54 | 1.00 | [6, 7] | Group20 | 4.84 | 3.00 |
| GO:0015399 | primary active transmembrane transporter activity | GO_BiologicalProcess-Custom-GOA_05.03.2019_00h00 | 0.47 | 1.00 | 0.54 | 1.00 | [7] | Group20 | 5.88 | 6.00 |
| GO:0022853 | active ion transmembrane transporter activity | GO_BiologicalProcess-Custom-GOA_05.03.2019_00h00 | 0.76 | 1.00 | 0.54 | 1.00 | [7] | Group20 | 4.84 | 3.00 |
| GO:0015405 | P-P-bond-hydrolysis-driven transmembrane transporter activity | GO_BiologicalProcess-Custom-GOA_05.03.2019_00h00 | 0.47 | 1.00 | 0.54 | 1.00 | [8] | Group20 | 5.88 | 6.00 |
| GO:0042625 | ATPase coupled ion transmembrane transporter activity | GO_BiologicalProcess-Custom-GOA_05.03.2019_00h00 | 0.76 | 1.00 | 0.54 | 1.00 | [7, 8, 10] | Group20 | 4.84 | 3.00 |

*Table S 3 List of significant GOterms 'biological process' for the top 5% of CNV measure.*

| GOID | GOTerm | Ontology Source | Term PValue | Term PValue Corrected with Bonferroni step down | Group PValue | Group PValue Corrected with Bonferroni step down | GOLevels | GOGroups | % Associated Genes | Nr. Genes |
|---|---|---|---|---|---|---|---|---|---|---|
| GO:0007031 | peroxisome organization | GO_BiologicalProcess-Custom-GOA_05.03.2019_00h00 | 0.00 | 0.19 | 0.00 | 0.04 | [4] | Group00 | 33.33 | 3.00 |
| GO:1901615 | organic hydroxy compound metabolic process | GO_BiologicalProcess-Custom-GOA_05.03.2019_00h00 | 0.05 | 1.00 | 0.05 | 0.70 | [3] | Group01 | 12.00 | 3.00 |
| GO:0043933 | protein-containing complex subunit organization | GO_BiologicalProcess-Custom-GOA_05.03.2019_00h00 | 0.24 | 1.00 | 0.24 | 1.00 | [3] | Group02 | 6.12 | 3.00 |
| GO:0044283 | small molecule biosynthetic process | GO_BiologicalProcess-Custom-GOA_05.03.2019_00h00 | 0.31 | 1.00 | 0.31 | 1.00 | [3] | Group03 | 5.00 | 6.00 |
| GO:0006629 | lipid metabolic process | GO_BiologicalProcess-Custom-GOA_05.03.2019_00h00 | 0.21 | 1.00 | 0.21 | 1.00 | [3] | Group04 | 5.60 | 7.00 |
| GO:0008610 | lipid biosynthetic process | GO_BiologicalProcess-Custom-GOA_05.03.2019_00h00 | 0.13 | 1.00 | 0.21 | 1.00 | [4] | Group04 | 7.02 | 4.00 |
| GO:0016651 | oxidoreductase activity, acting on NAD(P)H | GO_BiologicalProcess-Custom- | 0.44 | 1.00 | 0.44 | 1.00 | [4] | Group05 | 5.45 | 3.00 |

| | | | | | | | | | | |
|---|---|---|---|---|---|---|---|---|---|---|
| | | GOA_05.03.2019_00h00 | | | | | | | | |
| GO:0003959 | NADPH dehydrogenase activity | GO_BiologicalProcess-Custom-GOA_05.03.2019_00h00 | 0.01 | 0.70 | 0.44 | 1.00 | [5] | Group05 | 21.43 | 3.00 |
| GO:0016903 | oxidoreductase activity, acting on the aldehyde or oxo group of donors | GO_BiologicalProcess-Custom-GOA_05.03.2019_00h00 | 0.07 | 1.00 | 0.07 | 0.86 | [4] | Group06 | 10.71 | 3.00 |
| GO:0016620 | oxidoreductase activity, acting on the aldehyde or oxo group of donors, NAD or NADP as acceptor | GO_BiologicalProcess-Custom-GOA_05.03.2019_00h00 | 0.03 | 1.00 | 0.07 | 0.86 | [5] | Group06 | 15.00 | 3.00 |
| GO:0006259 | DNA metabolic process | GO_BiologicalProcess-Custom-GOA_05.03.2019_00h00 | 0.67 | 0.67 | 0.67 | 0.67 | [4, 5, 6] | Group07 | 4.00 | 7.00 |
| GO:0004536 | deoxyribonuclease activity | GO_BiologicalProcess-Custom-GOA_05.03.2019_00h00 | 0.17 | 1.00 | 0.67 | 0.67 | [5, 6, 7, 8] | Group07 | 7.32 | 3.00 |
| GO:0004520 | endodeoxyribonuclease activity | GO_BiologicalProcess-Custom-GOA_05.03.2019_00h00 | 0.12 | 1.00 | 0.67 | 0.67 | [6, 7, 8, 9] | Group07 | 8.57 | 3.00 |
| GO:0043765 | T/G mismatch-specific endonuclease activity | GO_BiologicalProcess-Custom-GOA_05.03.2019_00h00 | 0.10 | 1.00 | 0.67 | 0.67 | [7, 8, 9, 10] | Group07 | 9.38 | 3.00 |
| GO:0006520 | cellular amino acid metabolic process | GO_BiologicalProcess-Custom-GOA_05.03.2019_00h00 | 0.22 | 1.00 | 0.53 | 1.00 | [3, 4, 6] | Group08 | 5.38 | 7.00 |
| GO:0006518 | peptide metabolic process | GO_BiologicalProcess-Custom-GOA_05.03.2019_00h00 | 0.57 | 1.00 | 0.53 | 1.00 | [4, 5] | Group08 | 4.09 | 9.00 |
| GO:0043604 | amide biosynthetic process | GO_BiologicalProcess-Custom-GOA_05.03.2019_00h00 | 0.58 | 1.00 | 0.53 | 1.00 | [5] | Group08 | 4.00 | 9.00 |
| GO:0043038 | amino acid activation | GO_BiologicalProcess-Custom-GOA_05.03.2019_00h00 | 0.12 | 1.00 | 0.53 | 1.00 | [4, 5, 7] | Group08 | 8.57 | 3.00 |
| GO:1901605 | alpha-amino acid metabolic process | GO_BiologicalProcess-Custom-GOA_05.03.2019_00h00 | 0.45 | 1.00 | 0.53 | 1.00 | [4, 5, 7] | Group08 | 5.17 | 3.00 |
| GO:0043043 | peptide biosynthetic process | GO_BiologicalProcess-Custom-GOA_05.03.2019_00h00 | 0.56 | 1.00 | 0.53 | 1.00 | [5, 6] | Group08 | 4.19 | 9.00 |
| GO:0006412 | translation | GO_BiologicalProcess-Custom-GOA_05.03.2019_00h00 | 0.56 | 1.00 | 0.53 | 1.00 | [5, 6, 7] | Group08 | 4.25 | 9.00 |
| GO:0043039 | tRNA aminoacylation | GO_BiologicalProcess-Custom-GOA_05.03.2019_00h00 | 0.12 | 1.00 | 0.53 | 1.00 | [5, 6, 8, 9] | Group08 | 8.57 | 3.00 |
| GO:0006418 | tRNA aminoacylation for protein translation | GO_BiologicalProcess-Custom-GOA_05.03.2019_00h00 | 0.11 | 1.00 | 0.53 | 1.00 | [6, 7, 8, 9, 10] | Group08 | 8.82 | 3.00 |

| | | | | | | | | | | |
|---|---|---|---|---|---|---|---|---|---|---|
| GO:0006796 | phosphate-containing compound metabolic process | GO_BiologicalProcess-Custom-GOA_05.03.2019_00h00 | 0.39 | 1.00 | 0.25 | 1.00 | [4] | Group09 | 4.07 | 24.00 |
| GO:0043412 | macromolecule modification | GO_BiologicalProcess-Custom-GOA_05.03.2019_00h00 | 0.24 | 1.00 | 0.25 | 1.00 | [4] | Group09 | 4.24 | 28.00 |
| GO:0036211 | protein modification process | GO_BiologicalProcess-Custom-GOA_05.03.2019_00h00 | 0.23 | 1.00 | 0.25 | 1.00 | [4, 5] | Group09 | 4.27 | 27.00 |
| GO:0044267 | cellular protein metabolic process | GO_BiologicalProcess-Custom-GOA_05.03.2019_00h00 | 0.14 | 1.00 | 0.25 | 1.00 | [4, 5] | Group09 | 4.30 | 38.00 |
| GO:0016310 | phosphorylation | GO_BiologicalProcess-Custom-GOA_05.03.2019_00h00 | 0.18 | 1.00 | 0.25 | 1.00 | [5] | Group09 | 4.74 | 18.00 |
| GO:0006464 | cellular protein modification process | GO_BiologicalProcess-Custom-GOA_05.03.2019_00h00 | 0.23 | 1.00 | 0.25 | 1.00 | [5, 6] | Group09 | 4.27 | 27.00 |
| GO:0016301 | kinase activity | GO_BiologicalProcess-Custom-GOA_05.03.2019_00h00 | 0.13 | 1.00 | 0.25 | 1.00 | [6] | Group09 | 4.93 | 18.00 |
| GO:0006468 | protein phosphorylation | GO_BiologicalProcess-Custom-GOA_05.03.2019_00h00 | 0.07 | 1.00 | 0.25 | 1.00 | [6, 7] | Group09 | 5.33 | 16.00 |
| GO:0018193 | peptidyl-amino acid modification | GO_BiologicalProcess-Custom-GOA_05.03.2019_00h00 | 0.18 | 1.00 | 0.25 | 1.00 | [6, 7] | Group09 | 5.49 | 9.00 |
| GO:0004672 | protein kinase activity | GO_BiologicalProcess-Custom-GOA_05.03.2019_00h00 | 0.07 | 1.00 | 0.25 | 1.00 | [7, 8] | Group09 | 5.33 | 16.00 |
| GO:0018212 | peptidyl-tyrosine modification | GO_BiologicalProcess-Custom-GOA_05.03.2019_00h00 | 0.06 | 1.00 | 0.25 | 1.00 | [7, 8] | Group09 | 7.50 | 6.00 |
| GO:0018108 | peptidyl-tyrosine phosphorylation | GO_BiologicalProcess-Custom-GOA_05.03.2019_00h00 | 0.06 | 1.00 | 0.25 | 1.00 | [7, 8, 9] | Group09 | 7.50 | 6.00 |
| GO:0004674 | protein serine/threonine kinase activity | GO_BiologicalProcess-Custom-GOA_05.03.2019_00h00 | 0.54 | 1.00 | 0.25 | 1.00 | [8, 9] | Group09 | 4.15 | 8.00 |
| GO:0004713 | protein tyrosine kinase activity | GO_BiologicalProcess-Custom-GOA_05.03.2019_00h00 | 0.06 | 1.00 | 0.25 | 1.00 | [8, 9, 10] | Group09 | 7.50 | 6.00 |
| GO:0051649 | establishment of localization in cell | GO_BiologicalProcess-Custom-GOA_05.03.2019_00h00 | 0.29 | 1.00 | 0.24 | 1.00 | [3] | Group10 | 5.26 | 6.00 |
| GO:0070727 | cellular macromolecule localization | GO_BiologicalProcess-Custom-GOA_05.03.2019_00h00 | 0.18 | 1.00 | 0.24 | 1.00 | [3] | Group10 | 5.71 | 6.00 |
| GO:0045184 | establishment of protein localization | GO_BiologicalProcess-Custom-GOA_05.03.2019_00h00 | 0.65 | 1.00 | 0.24 | 1.00 | [3, 4] | Group10 | 4.03 | 6.00 |

| | | | | | | | | | | |
|---|---|---|---|---|---|---|---|---|---|---|
| GO:0046907 | intracellular transport | GO_BiologicalProcess-Custom-GOA_05.03.2019_00h00 | 0.29 | 1.00 | 0.24 | 1.00 | [3, 4] | Group10 | 5.26 | 6.00 |
| GO:0016192 | vesicle-mediated transport | GO_BiologicalProcess-Custom-GOA_05.03.2019_00h00 | 0.03 | 1.00 | 0.24 | 1.00 | [4] | Group10 | 8.57 | 6.00 |
| GO:0034613 | cellular protein localization | GO_BiologicalProcess-Custom-GOA_05.03.2019_00h00 | 0.18 | 1.00 | 0.24 | 1.00 | [4] | Group10 | 5.71 | 6.00 |
| GO:0071702 | organic substance transport | GO_BiologicalProcess-Custom-GOA_05.03.2019_00h00 | 0.29 | 1.00 | 0.24 | 1.00 | [4] | Group10 | 4.69 | 12.00 |
| GO:0071705 | nitrogen compound transport | GO_BiologicalProcess-Custom-GOA_05.03.2019_00h00 | 0.24 | 1.00 | 0.24 | 1.00 | [4] | Group10 | 4.85 | 10.00 |
| GO:0042886 | amide transport | GO_BiologicalProcess-Custom-GOA_05.03.2019_00h00 | 0.65 | 1.00 | 0.24 | 1.00 | [5] | Group10 | 4.03 | 6.00 |
| GO:0015031 | protein transport | GO_BiologicalProcess-Custom-GOA_05.03.2019_00h00 | 0.64 | 1.00 | 0.24 | 1.00 | [4, 5, 6, 7] | Group10 | 4.08 | 6.00 |
| GO:0015833 | peptide transport | GO_BiologicalProcess-Custom-GOA_05.03.2019_00h00 | 0.65 | 1.00 | 0.24 | 1.00 | [5, 6] | Group10 | 4.03 | 6.00 |
| GO:0006886 | intracellular protein transport | GO_BiologicalProcess-Custom-GOA_05.03.2019_00h00 | 0.17 | 1.00 | 0.24 | 1.00 | [4, 5, 6, 7, 8] | Group10 | 5.88 | 6.00 |
| GO:0006865 | amino acid transport | GO_BiologicalProcess-Custom-GOA_05.03.2019_00h00 | 0.09 | 1.00 | 0.24 | 1.00 | [5, 7, 8] | Group10 | 9.68 | 3.00 |
| GO:0003333 | amino acid transmembrane transport | GO_BiologicalProcess-Custom-GOA_05.03.2019_00h00 | 0.08 | 1.00 | 0.24 | 1.00 | [6, 7, 8, 9] | Group10 | 10.00 | 3.00 |
| GO:0015171 | amino acid transmembrane transporter activity | GO_BiologicalProcess-Custom-GOA_05.03.2019_00h00 | 0.08 | 1.00 | 0.24 | 1.00 | [7, 8, 9, 10] | Group10 | 10.00 | 3.00 |
| GO:0006518 | peptide metabolic process | GO_BiologicalProcess-Custom-GOA_05.03.2019_00h00 | 0.57 | 1.00 | 0.66 | 1.00 | [4, 5] | Group11 | 4.09 | 9.00 |
| GO:0034248 | regulation of cellular amide metabolic process | GO_BiologicalProcess-Custom-GOA_05.03.2019_00h00 | 0.11 | 1.00 | 0.66 | 1.00 | [4, 5] | Group11 | 7.55 | 4.00 |
| GO:0051246 | regulation of protein metabolic process | GO_BiologicalProcess-Custom-GOA_05.03.2019_00h00 | 0.33 | 1.00 | 0.66 | 1.00 | [4, 5] | Group11 | 4.76 | 10.00 |
| GO:0032268 | regulation of cellular protein metabolic process | GO_BiologicalProcess-Custom-GOA_05.03.2019_00h00 | 0.33 | 1.00 | 0.66 | 1.00 | [4, 5, 6] | Group11 | 4.78 | 10.00 |
| GO:0043604 | amide biosynthetic process | GO_BiologicalProcess-Custom-GOA_05.03.2019_00h00 | 0.58 | 1.00 | 0.66 | 1.00 | [5] | Group11 | 4.00 | 9.00 |
| GO:0043038 | amino acid activation | GO_BiologicalProcess-Custom- | 0.12 | 1.00 | 0.66 | 1.00 | [4, 5, 7] | Group11 | 8.57 | 3.00 |

| | | | | | | | | | | |
|---|---|---|---|---|---|---|---|---|---|---|
| | | GOA_05.03.2019_00h00 | | | | | | | | |
| GO:0010608 | posttranscriptional regulation of gene expression | GO_BiologicalProcess-Custom-GOA_05.03.2019_00h00 | 0.12 | 1.00 | 0.66 | 1.00 | [5, 6] | Group11 | 7.41 | 4.00 |
| GO:0043043 | peptide biosynthetic process | GO_BiologicalProcess-Custom-GOA_05.03.2019_00h00 | 0.56 | 1.00 | 0.66 | 1.00 | [5, 6] | Group11 | 4.19 | 9.00 |
| GO:0006412 | translation | GO_BiologicalProcess-Custom-GOA_05.03.2019_00h00 | 0.56 | 1.00 | 0.66 | 1.00 | [5, 6, 7] | Group11 | 4.25 | 9.00 |
| GO:0006413 | translational initiation | GO_BiologicalProcess-Custom-GOA_05.03.2019_00h00 | 0.14 | 1.00 | 0.66 | 1.00 | [3, 6, 7, 8] | Group11 | 8.11 | 3.00 |
| GO:0006417 | regulation of translation | GO_BiologicalProcess-Custom-GOA_05.03.2019_00h00 | 0.11 | 1.00 | 0.66 | 1.00 | [5, 6, 7, 8] | Group11 | 7.55 | 4.00 |
| GO:0043039 | tRNA aminoacylation | GO_BiologicalProcess-Custom-GOA_05.03.2019_00h00 | 0.12 | 1.00 | 0.66 | 1.00 | [5, 6, 8, 9] | Group11 | 8.57 | 3.00 |
| GO:0045182 | translation regulator activity | GO_BiologicalProcess-Custom-GOA_05.03.2019_00h00 | 0.09 | 1.00 | 0.66 | 1.00 | [6, 7, 8, 9] | Group11 | 8.16 | 4.00 |
| GO:0006418 | tRNA aminoacylation for protein translation | GO_BiologicalProcess-Custom-GOA_05.03.2019_00h00 | 0.11 | 1.00 | 0.66 | 1.00 | [6, 7, 8, 9, 10] | Group11 | 8.82 | 3.00 |
| GO:0008135 | translation factor activity, RNA binding | GO_BiologicalProcess-Custom-GOA_05.03.2019_00h00 | 0.03 | 1.00 | 0.66 | 1.00 | [6, 7, 8, 9, 10, 11] | Group11 | 11.76 | 4.00 |
| GO:0090079 | translation regulator activity, nucleic acid binding | GO_BiologicalProcess-Custom-GOA_05.03.2019_00h00 | 0.03 | 1.00 | 0.66 | 1.00 | [7, 8, 9, 10] | Group11 | 11.76 | 4.00 |
| GO:0003743 | translation initiation factor activity | GO_BiologicalProcess-Custom-GOA_05.03.2019_00h00 | 0.03 | 1.00 | 0.66 | 1.00 | [4, 7, 8, 9, 10, 11, 12] | Group11 | 15.00 | 3.00 |
| GO:0034248 | regulation of cellular amide metabolic process | GO_BiologicalProcess-Custom-GOA_05.03.2019_00h00 | 0.11 | 1.00 | 0.33 | 1.00 | [4, 5] | Group12 | 7.55 | 4.00 |
| GO:0051246 | regulation of protein metabolic process | GO_BiologicalProcess-Custom-GOA_05.03.2019_00h00 | 0.33 | 1.00 | 0.33 | 1.00 | [4, 5] | Group12 | 4.76 | 10.00 |
| GO:0032268 | regulation of cellular protein metabolic process | GO_BiologicalProcess-Custom-GOA_05.03.2019_00h00 | 0.33 | 1.00 | 0.33 | 1.00 | [4, 5, 6] | Group12 | 4.78 | 10.00 |
| GO:0051348 | negative regulation of transferase activity | GO_BiologicalProcess-Custom-GOA_05.03.2019_00h00 | 0.62 | 1.00 | 0.33 | 1.00 | [5] | Group12 | 4.03 | 5.00 |
| GO:0010608 | posttranscriptional regulation of gene expression | GO_BiologicalProcess-Custom-GOA_05.03.2019_00h00 | 0.12 | 1.00 | 0.33 | 1.00 | [5, 6] | Group12 | 7.41 | 4.00 |
| GO:0006413 | translational initiation | GO_BiologicalProcess-Custom-GOA_05.03.2019_00h00 | 0.14 | 1.00 | 0.33 | 1.00 | [3, 6, 7, 8] | Group12 | 8.11 | 3.00 |

| | | | | | | | | | | |
|---|---|---|---|---|---|---|---|---|---|---|
| GO:0006417 | regulation of translation | GO_BiologicalProcess-Custom-GOA_05.03.2019_00h00 | 0.11 | 1.00 | 0.33 | 1.00 | [5, 6, 7, 8] | Group12 | 7.55 | 4.00 |
| GO:0031400 | negative regulation of protein modification process | GO_BiologicalProcess-Custom-GOA_05.03.2019_00h00 | 0.62 | 1.00 | 0.33 | 1.00 | [5, 6, 7, 8] | Group12 | 4.00 | 5.00 |
| GO:0070647 | protein modification by small protein conjugation or removal | GO_BiologicalProcess-Custom-GOA_05.03.2019_00h00 | 0.61 | 1.00 | 0.33 | 1.00 | [6, 7] | Group12 | 4.20 | 5.00 |
| GO:1903320 | regulation of protein modification by small protein conjugation or removal | GO_BiologicalProcess-Custom-GOA_05.03.2019_00h00 | 0.61 | 1.00 | 0.33 | 1.00 | [6, 7, 8] | Group12 | 4.24 | 5.00 |
| GO:0032446 | protein modification by small protein conjugation | GO_BiologicalProcess-Custom-GOA_05.03.2019_00h00 | 0.61 | 1.00 | 0.33 | 1.00 | [7, 8] | Group12 | 4.20 | 5.00 |
| GO:0045182 | translation regulator activity | GO_BiologicalProcess-Custom-GOA_05.03.2019_00h00 | 0.09 | 1.00 | 0.33 | 1.00 | [6, 7, 8, 9] | Group12 | 8.16 | 4.00 |
| GO:1903321 | negative regulation of protein modification by small protein conjugation or removal | GO_BiologicalProcess-Custom-GOA_05.03.2019_00h00 | 0.61 | 1.00 | 0.33 | 1.00 | [6, 7, 8, 9] | Group12 | 4.24 | 5.00 |
| GO:1903322 | positive regulation of protein modification by small protein conjugation or removal | GO_BiologicalProcess-Custom-GOA_05.03.2019_00h00 | 0.61 | 1.00 | 0.33 | 1.00 | [6, 7, 8, 9] | Group12 | 4.24 | 5.00 |
| GO:0008135 | translation factor activity, RNA binding | GO_BiologicalProcess-Custom-GOA_05.03.2019_00h00 | 0.03 | 1.00 | 0.33 | 1.00 | [6, 7, 8, 9, 10, 11] | Group12 | 11.76 | 4.00 |
| GO:0016567 | protein ubiquitination | GO_BiologicalProcess-Custom-GOA_05.03.2019_00h00 | 0.61 | 1.00 | 0.33 | 1.00 | [8, 9] | Group12 | 4.20 | 5.00 |
| GO:0031396 | regulation of protein ubiquitination | GO_BiologicalProcess-Custom-GOA_05.03.2019_00h00 | 0.61 | 1.00 | 0.33 | 1.00 | [7, 8, 9, 10] | Group12 | 4.24 | 5.00 |
| GO:0090079 | translation regulator activity, nucleic acid binding | GO_BiologicalProcess-Custom-GOA_05.03.2019_00h00 | 0.03 | 1.00 | 0.33 | 1.00 | [7, 8, 9, 10] | Group12 | 11.76 | 4.00 |
| GO:0051438 | regulation of ubiquitin-protein transferase activity | GO_BiologicalProcess-Custom-GOA_05.03.2019_00h00 | 0.61 | 1.00 | 0.33 | 1.00 | [5, 8, 9, 10, 11] | Group12 | 4.24 | 5.00 |
| GO:0003743 | translation initiation factor activity | GO_BiologicalProcess-Custom-GOA_05.03.2019_00h00 | 0.03 | 1.00 | 0.33 | 1.00 | [4, 7, 8, 9, 10, 11, 12] | Group12 | 15.00 | 3.00 |
| GO:0031397 | negative regulation of protein ubiquitination | GO_BiologicalProcess-Custom-GOA_05.03.2019_00h00 | 0.61 | 1.00 | 0.33 | 1.00 | [7, 8, 9, 10, 11] | Group12 | 4.24 | 5.00 |

| | | | | | | | | | | |
|---|---|---|---|---|---|---|---|---|---|---|
| GO:0031398 | positive regulation of protein ubiquitination | GO_BiologicalProcess-Custom-GOA_05.03.2019_00h00 | 0.61 | 1.00 | 0.33 | 1.00 | [7, 8, 9, 10, 11] | Group12 | 4.24 | 5.00 |
| GO:0051443 | positive regulation of ubiquitin-protein transferase activity | GO_BiologicalProcess-Custom-GOA_05.03.2019_00h00 | 0.61 | 1.00 | 0.33 | 1.00 | [6, 8, 9, 10, 11, 12] | Group12 | 4.24 | 5.00 |
| GO:0051444 | negative regulation of ubiquitin-protein transferase activity | GO_BiologicalProcess-Custom-GOA_05.03.2019_00h00 | 0.61 | 1.00 | 0.33 | 1.00 | [6, 8, 9, 10, 11, 12] | Group12 | 4.24 | 5.00 |
| GO:0042493 | response to drug | GO_BiologicalProcess-Custom-GOA_05.03.2019_00h00 | 0.23 | 1.00 | 0.48 | 1.00 | [3] | Group13 | 6.25 | 3.00 |
| GO:0051649 | establishment of localization in cell | GO_BiologicalProcess-Custom-GOA_05.03.2019_00h00 | 0.29 | 1.00 | 0.48 | 1.00 | [3] | Group13 | 5.26 | 6.00 |
| GO:0070727 | cellular macromolecule localization | GO_BiologicalProcess-Custom-GOA_05.03.2019_00h00 | 0.18 | 1.00 | 0.48 | 1.00 | [3] | Group13 | 5.71 | 6.00 |
| GO:0045184 | establishment of protein localization | GO_BiologicalProcess-Custom-GOA_05.03.2019_00h00 | 0.65 | 1.00 | 0.48 | 1.00 | [3, 4] | Group13 | 4.03 | 6.00 |
| GO:0046907 | intracellular transport | GO_BiologicalProcess-Custom-GOA_05.03.2019_00h00 | 0.29 | 1.00 | 0.48 | 1.00 | [3, 4] | Group13 | 5.26 | 6.00 |
| GO:0015893 | drug transport | GO_BiologicalProcess-Custom-GOA_05.03.2019_00h00 | 0.23 | 1.00 | 0.48 | 1.00 | [4] | Group13 | 6.25 | 3.00 |
| GO:0034613 | cellular protein localization | GO_BiologicalProcess-Custom-GOA_05.03.2019_00h00 | 0.18 | 1.00 | 0.48 | 1.00 | [4] | Group13 | 5.71 | 6.00 |
| GO:0055085 | transmembrane transport | GO_BiologicalProcess-Custom-GOA_05.03.2019_00h00 | 0.29 | 1.00 | 0.48 | 1.00 | [4] | Group13 | 4.60 | 12.00 |
| GO:0071702 | organic substance transport | GO_BiologicalProcess-Custom-GOA_05.03.2019_00h00 | 0.29 | 1.00 | 0.48 | 1.00 | [4] | Group13 | 4.69 | 12.00 |
| GO:0071705 | nitrogen compound transport | GO_BiologicalProcess-Custom-GOA_05.03.2019_00h00 | 0.24 | 1.00 | 0.48 | 1.00 | [4] | Group13 | 4.85 | 10.00 |
| GO:0006820 | anion transport | GO_BiologicalProcess-Custom-GOA_05.03.2019_00h00 | 0.11 | 1.00 | 0.48 | 1.00 | [5] | Group13 | 6.85 | 5.00 |
| GO:0006855 | drug transmembrane transport | GO_BiologicalProcess-Custom-GOA_05.03.2019_00h00 | 0.23 | 1.00 | 0.48 | 1.00 | [5] | Group13 | 6.25 | 3.00 |
| GO:0015849 | organic acid transport | GO_BiologicalProcess-Custom-GOA_05.03.2019_00h00 | 0.03 | 1.00 | 0.48 | 1.00 | [5] | Group13 | 12.12 | 4.00 |
| GO:0022857 | transmembrane transporter activity | GO_BiologicalProcess-Custom-GOA_05.03.2019_00h00 | 0.48 | 1.00 | 0.48 | 1.00 | [5] | Group13 | 4.23 | 11.00 |

| | | | | | | | | | | |
|---|---|---|---|---|---|---|---|---|---|---|
| GO:0042886 | amide transport | GO_BiologicalProcess-Custom-GOA_05.03.2019_00h00 | 0.65 | 1.00 | 0.48 | 1.00 | [5] | Group13 | 4.03 | 6.00 |
| GO:0015031 | protein transport | GO_BiologicalProcess-Custom-GOA_05.03.2019_00h00 | 0.64 | 1.00 | 0.48 | 1.00 | [4, 5, 6, 7] | Group13 | 4.08 | 6.00 |
| GO:0015711 | organic anion transport | GO_BiologicalProcess-Custom-GOA_05.03.2019_00h00 | 0.04 | 1.00 | 0.48 | 1.00 | [5, 6] | Group13 | 9.09 | 5.00 |
| GO:0015833 | peptide transport | GO_BiologicalProcess-Custom-GOA_05.03.2019_00h00 | 0.65 | 1.00 | 0.48 | 1.00 | [5, 6] | Group13 | 4.03 | 6.00 |
| GO:1903825 | organic acid transmembrane transport | GO_BiologicalProcess-Custom-GOA_05.03.2019_00h00 | 0.02 | 1.00 | 0.48 | 1.00 | [5, 6] | Group13 | 12.50 | 4.00 |
| GO:0006886 | intracellular protein transport | GO_BiologicalProcess-Custom-GOA_05.03.2019_00h00 | 0.17 | 1.00 | 0.48 | 1.00 | [4, 5, 6, 7, 8] | Group13 | 5.88 | 6.00 |
| GO:0015238 | drug transmembrane transporter activity | GO_BiologicalProcess-Custom-GOA_05.03.2019_00h00 | 0.23 | 1.00 | 0.48 | 1.00 | [6] | Group13 | 6.25 | 3.00 |
| GO:0015318 | inorganic molecular entity transmembrane transporter activity | GO_BiologicalProcess-Custom-GOA_05.03.2019_00h00 | 0.39 | 1.00 | 0.48 | 1.00 | [6] | Group13 | 4.71 | 8.00 |
| GO:0030001 | metal ion transport | GO_BiologicalProcess-Custom-GOA_05.03.2019_00h00 | 0.19 | 1.00 | 0.48 | 1.00 | [6] | Group13 | 6.98 | 3.00 |
| GO:0098656 | anion transmembrane transport | GO_BiologicalProcess-Custom-GOA_05.03.2019_00h00 | 0.07 | 1.00 | 0.48 | 1.00 | [6] | Group13 | 7.81 | 5.00 |
| GO:0005342 | organic acid transmembrane transporter activity | GO_BiologicalProcess-Custom-GOA_05.03.2019_00h00 | 0.02 | 1.00 | 0.48 | 1.00 | [6, 7] | Group13 | 12.50 | 4.00 |
| GO:0046942 | carboxylic acid transport | GO_BiologicalProcess-Custom-GOA_05.03.2019_00h00 | 0.03 | 1.00 | 0.48 | 1.00 | [6, 7] | Group13 | 12.12 | 4.00 |
| GO:0006865 | amino acid transport | GO_BiologicalProcess-Custom-GOA_05.03.2019_00h00 | 0.09 | 1.00 | 0.48 | 1.00 | [5, 7, 8] | Group13 | 9.68 | 3.00 |
| GO:0008509 | anion transmembrane transporter activity | GO_BiologicalProcess-Custom-GOA_05.03.2019_00h00 | 0.07 | 1.00 | 0.48 | 1.00 | [7] | Group13 | 7.81 | 5.00 |
| GO:0008514 | organic anion transmembrane transporter activity | GO_BiologicalProcess-Custom-GOA_05.03.2019_00h00 | 0.03 | 1.00 | 0.48 | 1.00 | [6, 7, 8] | Group13 | 10.00 | 5.00 |
| GO:1905039 | carboxylic acid transmembrane transport | GO_BiologicalProcess-Custom-GOA_05.03.2019_00h00 | 0.02 | 1.00 | 0.48 | 1.00 | [6, 7, 8] | Group13 | 12.50 | 4.00 |
| GO:0003333 | amino acid transmembrane transport | GO_BiologicalProcess-Custom-GOA_05.03.2019_00h00 | 0.08 | 1.00 | 0.48 | 1.00 | [6, 7, 8, 9] | Group13 | 10.00 | 3.00 |

| | | | | | | | | | | |
|---|---|---|---|---|---|---|---|---|---|---|
| GO:0046873 | metal ion transmembrane transporter activity | GO_BiologicalProcess-Custom-GOA_05.03.2019_00h00 | 0.10 | 1.00 | 0.48 | 1.00 | [7, 8, 9] | Group13 | 9.38 | 3.00 |
| GO:0046943 | carboxylic acid transmembrane transporter activity | GO_BiologicalProcess-Custom-GOA_05.03.2019_00h00 | 0.02 | 1.00 | 0.48 | 1.00 | [7, 8, 9] | Group13 | 12.50 | 4.00 |
| GO:0015171 | amino acid transmembrane transporter activity | GO_BiologicalProcess-Custom-GOA_05.03.2019_00h00 | 0.08 | 1.00 | 0.48 | 1.00 | [7, 8, 9, 10] | Group13 | 10.00 | 3.00 |

.

*Table S 4 List of gene annotations and Protein IDs for the top 5% of divergence measures $F_{ST}$, $d_{XY}$, and copy-number variation (CNV).*

| Heavy metal tolerance strategy | Strategy sub-category | Predicted/potential function (citation) | Annotation | CDS scaffold: start coordinate | statistic | Protein ID |
|---|---|---|---|---|---|---|
| **EXCLUSION /STORAGE:** | Metal ion transporters | Zinc transporter from cytosol into vesicle/vacuole Eide (2006) | Cation diffusion facilitator (CDF) transporter | 3:1754751 | FST | 2898571 |
| | | | | 3:1714851 | FST | 72605 |
| | | | | 3:1754751 | FST | 72657 |
| | | Iron transporter from cytosol into vesicle/vacuole (Ferrol et al. 2016) | Iron permease (FTR1) | 13:879566 | DXY | 2861746 |
| | Transport and transfer proteins, interacting with membranes | Vesicle transport (Brandizzi & Barlowe 2013) | ER-Golgi trafficking TRAPP I complex 85 kDa subunit | 1:2526703 | FST | 2760061 |
| | | Vesicle transport (Striegl et al. 2010; Tewari et al. 2010) | Armadillo-type fold | 2:1624106 | DXY | 2848533 |
| | | | | 3:386596 | CNV | 2764019 |
| | | | | 3:411345 | CNV | 2404130 |
| | | | | 4:1597784 | FST | 2852156 |

| | | | | | | |
|---|---|---|---|---|---|---|
| | | Vesicle transport (Conchon et al. 1999) | Vesicle transport protein, Got1/SFT2-like protein | 5:1537688 | CNV | 2767923 |
| | | Vesicle transport (Chen et al. 2017) | Exocyst complex component Sec10-like protein | 10:379697 | FST | 2858628 |
| | | Vesicle transport Lunkowitz (2011) | OPT oligopeptide transporter protein | 43:118643 | CNV | 2534791 |
| | | Zinc binding site (Zhang et al) | Rab geranylgeranyltransferase | 19:694943 | FST | 2921934 |
| | | Inhibit Rab geranylgeranyltransferase (Lackner et al) | Farnesyltranstransferase | 9:85745 | DXY | 27093 |
| | | | | 9:85745 | CNV | 27093 |
| **IMMOBILIZATION:** | Chelating agents within cytosol | Synthesis of Nicotianamine a cadmium chelating agent in cytosol (Aloui et al. 2009) | S-adenosyl-L-methionine-dependent methyltransferase | 2:1354115 | FST | 2848300 |
| | | | | 6:1123173 | CNV | 2854576 |
| | | | | 10:320100 | FST | 2858598 |
| | | Zinc chelating in cytosol (Lima et al. 1996) | HIT-like protein | 35:185000 | DXY | 2872095 |
| | Chelating agents outside the cell | Response to stress and chelating secreted protein (Ferrol et al. 2016) | Heat shock protein 70 family | 5:429146 | DXY | 2852980 |
| | | | | 15:533929 | DXY | 2751744 |
| | | | | 38:13252 | CNV | 2925132 |
| | | Metal ion chelating secreted protein (Ferrol et al. 2016) | Fungal hydrophobin-domain containing protein | 18:709254 | DXY | 2921647 |
| **ROS DETOXIFICATION:** | Oxidative stress relief | Antioxidant (Fang et al. 2002) | Manganese/iron superoxide dismutase | 1:3518896 | DXY | 2759231 |
| | | | | 36:176708 | DXY | 2470845 |
| | | Antioxidant (Johnson et al. 2003) | FAD-binding | 2:2438021 | DXY | 2849043 |
| | | | | 8:966734 | FST | 2856896 |
| | | | | 18:346289 | FST | 2864945 |
| | | | | 18:346289 | CNV | 2864945 |
| | | | | 19:804910 | DXY | 2902232 |
| | | | | 25:198508 | DXY | 2923258 |
| | | Antioxidant (Johnson et al. 2003) | Ferric reductase NAD binding domain | 7:1269395 | CNV | 2623337 |
| | | | | 7:1269395 | DXY | 2623337 |
| | | Antioxidant (Jozefczak et al. 2012) | Glutathione S-transferase | 8:354997 | FST | 2634392 |
| | | Pollution detoxification (Van den Brink et al) | Cytochrome P450 | 3:1019981 | DXY | 2898392 |
| | | | | 4:2311001 | DXY | 2765599 |
| | | | | 6:631643 | DXY | 2854272 |
| | | | | 7:314615 | DXY | 83884 |
| | | | | 12:39991 | FST | 2130781 |

| | | | | | | |
|---|---|---|---|---|---|---|
| | | | | 29:14823 | FST | 2950944 |
| | | | | 38:192801 | FST | 83946 |
| | | | | 40:55014 | FST | 2986292 |
| | | | | 5:459308 | FST | 12943 |
| | | | | 8:1236128 | FST | 24266 |
| | | | | 14:903209 | VST | 1888098 |
| | | | | 25:90970 | VST | 2902692 |
| | | | | 6:631999 | VST | 2854272 |

*Table S 5 Fungal isolates Short Read Archive codes, Illumina library information and ReadGroup codes.*

| Sample name | Short read archive code | Sample_id | ReadGroupID | RG_LB | RG_PL | RG_PU |
|---|---|---|---|---|---|---|
| UH-Slu-B10 | PRJNA456016 | 502931_1178750 | 12198.8.244335.CACTGAC-TGTCAGT | CHYAH | illumina | CACTGAC-TGTCAGT |
| UH-Slu-B12 | PRJNA456015 | 502931_1178751 | 12198.8.244335.TGCTTGG-ACCAAGC | CHYAN | illumina | TGCTTGG-ACCAAGC |
| UH-Slu-B2a | PRJNA456014 | 502931_1178744 | 12198.8.244335.GCCATAA-GTTATGG | CHXZY | illumina | GCCATAA-GTTATGG |
| UH-Slu-B3c | PRJNA456013 | 502931_1178745 | 12198.8.244335.ATGCCTG-ACAGGCA | CHXZZ | illumina | ATGCCTG-ACAGGCA |
| UH-Slu-B4a | PRJNA456012 | 502931_1178746 | 12198.8.244335.GAAGTAC-GGTACTT | CHYAA | illumina | GAAGTAC-GGTACTT |
| UH-Slu-B6 | PRJNA456011 | 502931_1178747 | 12198.8.244335.CGAACTG-ACAGTTC | CHYAB | illumina | CGAACTG-ACAGTTC |
| UH-Slu-B7b | PRJNA456010 | 502931_1178748 | 12198.8.244335.TAGTGAC-GGTCACT | CHYAC | illumina | TAGTGAC-GGTCACT |
| UH-Slu-B9a | PRJNA456009 | 502931_1178749 | 12198.8.244335.TGGCATG-ACATGCC | CHYAG | illumina | TGGCATG-ACATGCC |
| UH-Slu-DS16 | PRJNA459075 | 502931_1151576 | 12232.1.246284.ACCATCC-TGGATGG | BXUOO | illumina | ACCATCC-TGGATGG |
| UH-Slu-DS3 | PRJNA459306 | 502931_1151572 | 12232.1.246284.ACAGCAA-GTTGCTG | BXUOC | illumina | ACAGCAA-GTTGCTG |
| UH-Slu-DS4 | PRJNA456008 | 502931_1143066 | 12198.1.244228.GTGAGCT-AAGCTCA | CGNZX | illumina | GTGAGCT-AAGCTCA |
| UH-Slu-DS5 | PRJNA456007 | 502931_1143067 | 12198.4.244282.GAGCTCA-TTGAGCT | CGNXZ | illumina | GAGCTCA-TTGAGCT |
| UH-Slu-DS7 | PRJNA456006 | 502931_1143068 | 12198.4.244282.ACCATCC-TGGATGG | CGNZB | illumina | ACCATCC-TGGATGG |
| UH-Slu-DS9 | PRJNA459308 | 502931_1151574 | 12232.1.246284.TCATCAC-GGTGATG | BXUOH | illumina | TCATCAC-GGTGATG |
| UH-Slu-Ew1 | PRJNA456004 | 502931_1143059 | 12198.1.244228.CTGACAC-TGTGTCA | CGNYH | illumina | CTGACAC-TGTGTCA |
| UH-Slu-Ew2 | PRJNA456001 | 502931_1143060 | 12198.2.244246.GTTCGGT-AACCGAA | CGNYW | illumina | GTTCGGT-AACCGAA |
| UH-Slu-Ew3 | PRJNA459303 | 502931_1151569 | 12198.5.244301.GGCTAC | CGCZZ | illumina | GGCTAC |
| UH-Slu-Ew7 | PRJNA455999 | 502931_1143062 | 12198.3.244264.ACAGCAA-GTTGCTG | CGNYX | illumina | ACAGCAA-GTTGCTG |
| UH-Slu-LM12 | PRJNA455997 | 502931_1143052 | 12198.1.244228.TCCGAGT-AACTCGG | CGNZH | illumina | TCCGAGT-AACTCGG |
| UH-Slu-LM3 | PRJNA455995 | 502931_1143050 | 12198.2.244246.CAATCGA-GTCGATT | CGNZY | illumina | CAATCGA-GTCGATT |
| UH-Slu-LM5 | PRJNA455994 | 502931_1143051 | 12198.2.244246.GCCTTGT-AACAAGG | CGNYG | illumina | GCCTTGT-AACAAGG |
| UH-Slu-LmD7 | PRJNA455992 | 502931_1143078 | 12198.1.244228.TACCAAC-GGTTGGT | CGNYC | illumina | TACCAAC-GGTTGGT |
| UH-Slu-Sl1 | PRJNA459294 | 502931_1151559 | 12232.1.246284.GAGCTCA-TTGAGCT | BXUNH | illumina | GAGCTCA-TTGAGCT |

| | | | | | | |
|---|---|---|---|---|---|---|
| UH-Slu-Sl2 | PRJNA459295 | 502931_1151560 | 12232.1.246284.ATAGCGG-ACCGCTA | BXUNN | illumina | ATAGCGG-ACCGCTA |
| UH-Slu-Sl20 | PRJNA459296 | 502931_1151561 | 12232.1.246284.CGGTTGT-AACAACC | BXUNO | illumina | CGGTTGT-AACAACC |
| UH-Slu-Sl22 | PRJNA459297 | 502931_1151562 | 12232.1.246284.TACCAAC-GGTTGGT | BXUNP | illumina | TACCAAC-GGTTGGT |
| UH-Slu-Sl24 | PRJNA459298 | 502931_1151564 | 12232.1.246284.CTGACAC-TGTGTCA | BXUNT | illumina | CTGACAC-TGTGTCA |
| UH-Slu-N3 | PRJNA459079 | 502931_1151580 | 12232.1.246284.AGTCTCA-GTGAGAC | BXUOU | illumina | AGTCTCA-GTGAGAC |
| UH-Slu-N4 | PRJNA459080 | 502931_1151581 | 12232.1.246284.CCTCAGT-AACTGAG | BXUOW | illumina | CCTCAGT-AACTGAG |
| UH-Slu-N5 | PRJNA459081 | 502931_1151582 | 12232.1.246284.TTCGTAC-GGTACGA | BXUOX | illumina | TTCGTAC-GGTACGA |
| UH-Slu-Na2 | PRJNA455990 | 502931_1143070 | 12198.4.244282.ATAGCGG-ACCGCTA | CGNYA | illumina | ATAGCGG-ACCGCTA |
| UH-Slu-Na5 | PRJNA455987 | 502931_1143073 | 12198.2.244246.TCATCAC-GGTGATG | CGNYZ | illumina | TCATCAC-GGTGATG |
| UH-Slu-P2 | PRJNA455983 | 502931_1143054 | 12198.4.244282.TCGCTGT-AACAGCG | CGNYP | illumina | TCGCTGT-AACAGCG |
| UH-Slu-P8 | PRJNA455981 | 502931_1143056 | 12198.4.244282.GCTACGT-AACGTAG | CGNZC | illumina | GCTACGT-AACGTAG |
| UH-Slu-PD13 | PRJNA455980 | 502931_1143075 | 12198.3.244264.TGTACAC-GGTGTAC | CGNYO | illumina | TGTACAC-GGTGTAC |
| UH-Slu-PD14 | PRJNA455979 | 502931_1143076 | 12198.2.244246.GGACTGT-AACAGTC | CGNYT | illumina | GGACTGT-AACAGTC |
| UH-Slu-Sl31 | PRJNA459300 | 502931_1151566 | 12232.1.246284.TGTACAC-GGTGTAC | BXUNW | illumina | TGTACAC-GGTGTAC |
| UH-Slu-Sl33 | PRJNA459301 | 502931_1151567 | 12232.1.246284.TCGCTGT-AACAGCG | BXUNX | illumina | TCGCTGT-AACAGCG |
